# Strain-Programmable Patch for Diabetic Wound Healing

**DOI:** 10.1101/2021.06.07.447423

**Authors:** Georgios Theocharidis, Hyunwoo Yuk, Heejung Roh, Liu Wang, Ikram Mezghani, Jingjing Wu, Antonios Kafanas, Lihong Chen, Chuan Fei Guo, Navin Jayaswal, Xanthi-Leda Katopodi, Christoph S. Nabzdyk, Ioannis S. Vlachos, Aristidis Veves, Xuanhe Zhao

## Abstract

Chronic wounds with impaired healing capability such as diabetic foot ulcers (DFU) are devastating complications in diabetic patients, inflicting rapidly growing clinical and economic burdens in aging societies. Despite recent advances in therapeutic approaches, limited benefits of the existing solutions highlight the critical need for novel therapeutic solutions for diabetic wound healing. Here we propose a strain-programmable patch capable of rapid robust adhesion on and programmable mechanical contraction of wet wounded tissues over days to offer a new therapeutic platform for diabetic wounds. The strain-programmable patch, consisting of a dried bioadhesive layer and a pre-stretched elastomer backing, implements a hydration-based shape-memory mechanism to achieve both uniaxial and biaxial contractions and stress remodeling of wet wounds in a programmable manner. We develop theoretical and numerical models to rationally guide the strain-programming and mechanical modulation of wounds. *In vivo* rodent and *ex vivo* human skin culture models validate the programmability and efficacy of the proposed platform and identify mechanisms of action for accelerated diabetic wound healing.

**One Sentence Summary:** A strain-programmable bioadhesive patch is developed for accelerated closure and healing of wounds in diabetic mice and human skin.

## INTRODUCTION

Impaired wound healing capability and consequent chronic wounds such as diabetic foot ulcers (DFU) are one of the major and rapid growing complications in diabetic patients with over 750,000 new DFU each year (*1, 2*). Chronic DFU inflict significant clinical and economic burdens including 70,000 lower extremity amputations, a dramatic reduction in life quality (*2*) and associated costs over 11 billion dollars annually in the U.S. alone (*3*). Although various therapeutic strategies, such as bioengineered skin (*4, 5*) and growth factor-based treatments (*6*) have been introduced in clinical practice in the last few decades, their benefits are rather limited as more than 50% of treated DFU patients fail to respond (*7, 8*). The rapidly rising number of diabetic patients worldwide and the lack of effective treatment highlight the critical importance of developing new therapeutic solutions for diabetic wound healing.

Mechanical modulation of wounded or scarred skin has been a promising strategy to repair and remodel the skin in both animal models and human clinical trials (*9-17*). In addition, animal studies have indicated that the reduced contractibility of diabetic wounds compared to non-diabetic wounds is one of the sources of impaired diabetic wound healing (*7, 15, 18-20*). Therefore, mechanical modulation such as inducing contraction of diabetic wounds can be an attractive approach to accelerate diabetic wound healing. However, the potential therapeutic benefits of the mechanical modulation approach have not been well investigated for diabetic wounds such as DFU due to several technical limitations. Existing wound dressings and bandages for mechanical reinforcement or stimulation lack the capabilities to form rapid and robust adhesion on wet wounded skin over the long term (e.g., days) or to precisely program the mechanical contraction for wounds, limiting their use only for passive coverage, ineffective and uncontrolled contraction, and/or fully closed wounds (*13, 21-23*). To the best of our knowledge, there exists no method that can exert precisely-controlled and long-term contraction on wet wounded skin, leaving this potential therapeutic strategy untapped for diabetic wounds.

In this work, we report a strain-programmable patch capable of programmable and consistent mechanical contraction of wet wounded tissues over days as a promising therapeutic platform for accelerated healing of diabetic wounds. The strain-programmable patch synergistically combines a dry-crosslinking mechanism and a hydration-based shape-memory mechanism to simultaneously achieve robust, long-lasting, and on-demand detachable adhesion on wet wounded tissues and precisely-controlled mechanical modulation of the wounds, respectively. The strain-programmable patch takes the form of a thin flexible film consisting of a dry bioadhesive in the glassy state (high Young’s modulus) and a pre-stretched non-adhesive elastomer backing in the rubbery state (low Young’s modulus) with the programmed strain (Fig. 1A). The programmed strain in the patch is maintained due to the constraint of the dry bioadhesive. Once hydrated by the native physiological fluids and/or moisture from the wet wounded tissue, the bioadhesive rapidly and robustly adheres to the underlying tissue (fig. S1) and transits into a soft hydrogel in the rubbery state. Since the bioadhesive in the rubber state loses its ability to constrain the elastic recovery of the patch, the programmed strain in the patch is released to provide precisely-controlled mechanical contraction on the adhered wounded tissue (Fig. 1A). We perform systematic characterizations for the mechanical properties and adhesion performance of the strain-programmable patch and develop theoretical and numerical models to establish design principles for the strain-programming and mechanical modulation of wounds. The biocompatibility of the strain-programmable patch is investigated based on *in vitro* and *in vivo* rat models. We further validate the efficacy of the proposed patch for programmed contraction of wounded tissues and accelerated healing of diabetic wounds based on *in vivo* diabetic mice and *ex vivo* human skin culture models.

**Fig. 1.**
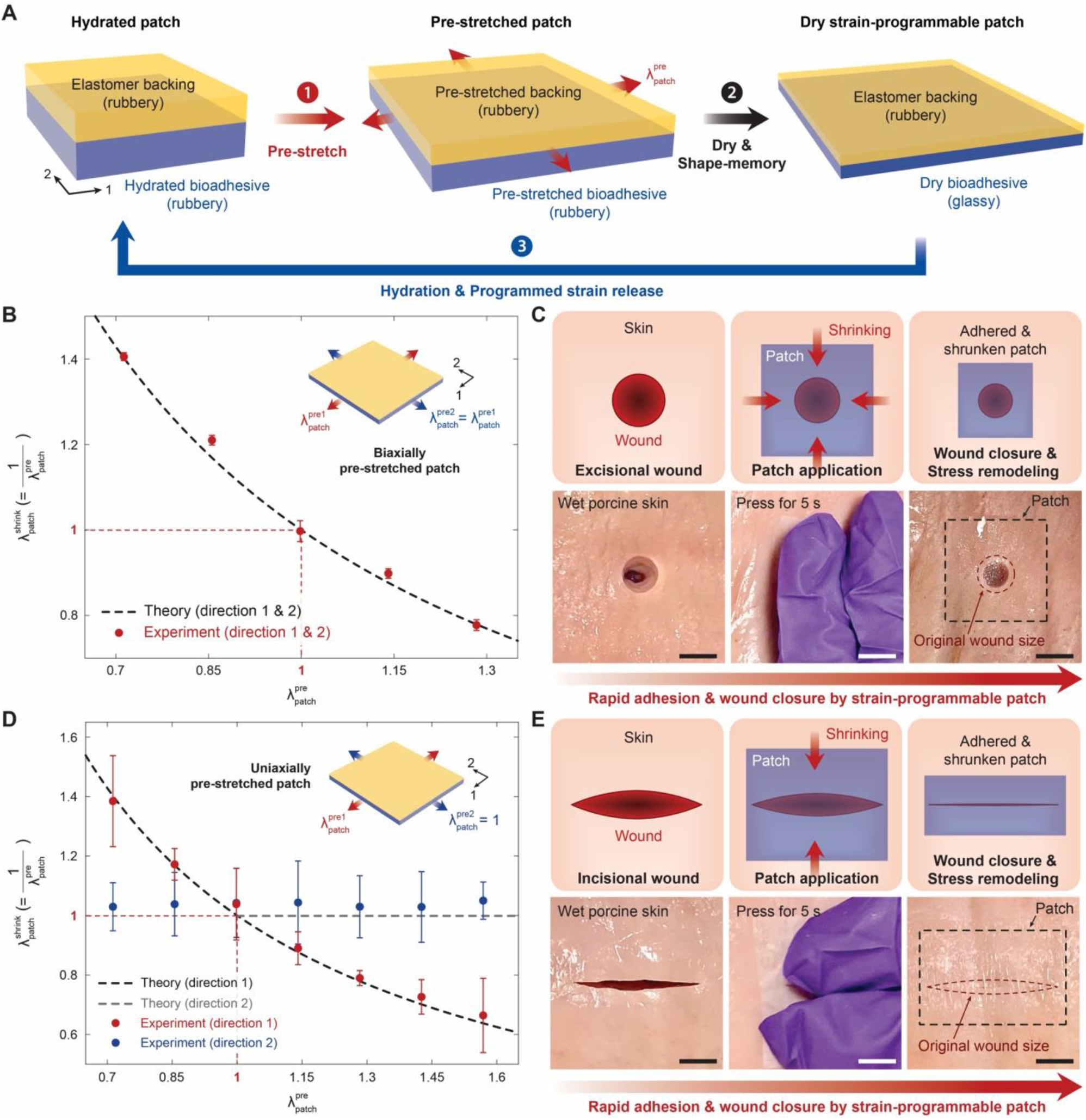
Design and mechanism of the strain-programmable patch. (**A**) Strain programming of the patch by a hydration-based shape-memory mechanism. (**B**) Theoretical and experimental values of 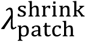 vs. 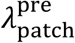 for isotropically strain-programmed patches. (**C**) Closure of a circular wound in a porcine skin by an isotropically strain-programmed patch. (**D**) Theoretical and experimental values of 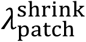 vs. 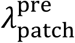 for anisotropically strain-programmed patches. (**E**) Closure of an incisional wound in a porcine skin by an anisotropically strain-programmed patch. Values in B and D represent the means ± SD (*n* = 4). Scale bars, 10 mm (C and E).

## RESULTS

### Design and mechanisms of the strain-programmable patch

The strain-programmable patch consists of two layers: (i) a non-adhesive elastomer backing based on a hydrophilic thermoplastic polyurethane and (ii) a bioadhesive layer based on crosslinked networks of poly(acrylic acid) grafted with *N*-hydroxysuccinimide ester (PAA-NHS ester) and chitosan (Fig. 1A). The hydration-based shape-memory mechanism of the strain-programmable patch relies on a drastic change in the mechanical properties of the bioadhesive layer based on its hydration states (*24-26*) (see Supplementary Text for details on the hydration-based shape-memory mechanism). The hydrated bioadhesive in the rubbery state is soft (i.e., Young’s modulus ∼ 40 kPa) and stretchable (i.e., over 4 times of the original length), whereas the dry bioadhesive becomes a glassy polymer with over 5 orders of magnitudes increase in stiffness (i.e., Young’s modulus ∼ 5 GPa) (fig. S2). To program the strain in the patch, an assembly of the hydrated bioadhesive layer bonded with the elastomer backing is pre-stretched along with two in-plane directions (i.e., directions 1 and 2 in Fig. 1A) by ratios of 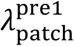 and 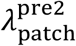, respectively. The pre-stretches on the assembly are maintained until the bioadhesive layer is dried to the glassy state. The glassy bioadhesive layer “freezes” the applied pre-stretches in itself (*27-29*) and constrains the elastomer backing from releasing the pre-stretches, due to the much higher rigidity of the glassy bioadhesive than the backing (Fig 1A and figs. S3 to S5; see Supplementary Text for details on the fabrication and strain-programming process).

Upon application of the strain-programmable patch on wet wounded tissues, the bioadhesive layer in the patch provides rapid robust adhesion to the wet wounded tissue surface within 5 s by absorbing the interfacial water between the patch and the tissue and forming crosslinks via the dry-crosslinking mechanism (*23, 26*) (fig. S1). Meanwhile, as the bioadhesive layer becomes hydrated, it quickly returns to the soft rubbery state within 30 s, during which the strain-programmable patch shrinks along with the two in-plane directions by ratios of 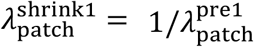 and 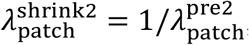, respectively (Fig. 1, B and D). This synergistic combination of the dry-crosslinking mechanism for rapid robust wet adhesion and the hydration-based shape-memory mechanism for strain-programming enables facile and highly effective mechanical modulation of wet wounded tissues by the strain-programmable patch (fig. S6). Notably, the biaxial programmability of the strain-programmable patch endows it with broad applicability to various types of wounds including isotropic open wounds (i.e., 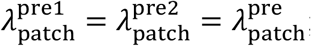; Fig. 1C and movie S1) as well as anisotropic incisional wounds (i.e., 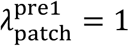; Fig. 1E and movie S2).

### Mechanical properties

The dry strain-programmable patch takes the form of a flexible thin dressing that can be applied to wet wounded tissues without any preparation process (Fig. 2, A and C). After adhering on tissues and releasing programmed strains, the swollen strain-programmable patch becomes a thin hydrogel layer with the tissue-like softness (Young’s modulus ∼ 50 kPa), stretchability (over 3.5 times of the original length) (Fig. 2, B and D), and high fracture toughness (over 400 J m^-2^) (fig. S7). Owing to the high programmability of the hydration-based shape-memory process, the contractile mechanical stresses generated by the strain-programmable patch can be precisely controlled based on the applied pre-stretches and the mechanical properties of the patch (Fig. 2, E and F; see Supplementary Text for details on the theoretical analysis). In particular, the mechanical properties and generated contractile stresses of the strain-programmable patch can be tuned by choosing various non-adhesive elastomer backing materials with different mechanical properties (fig. S8).

**Fig. 2.**
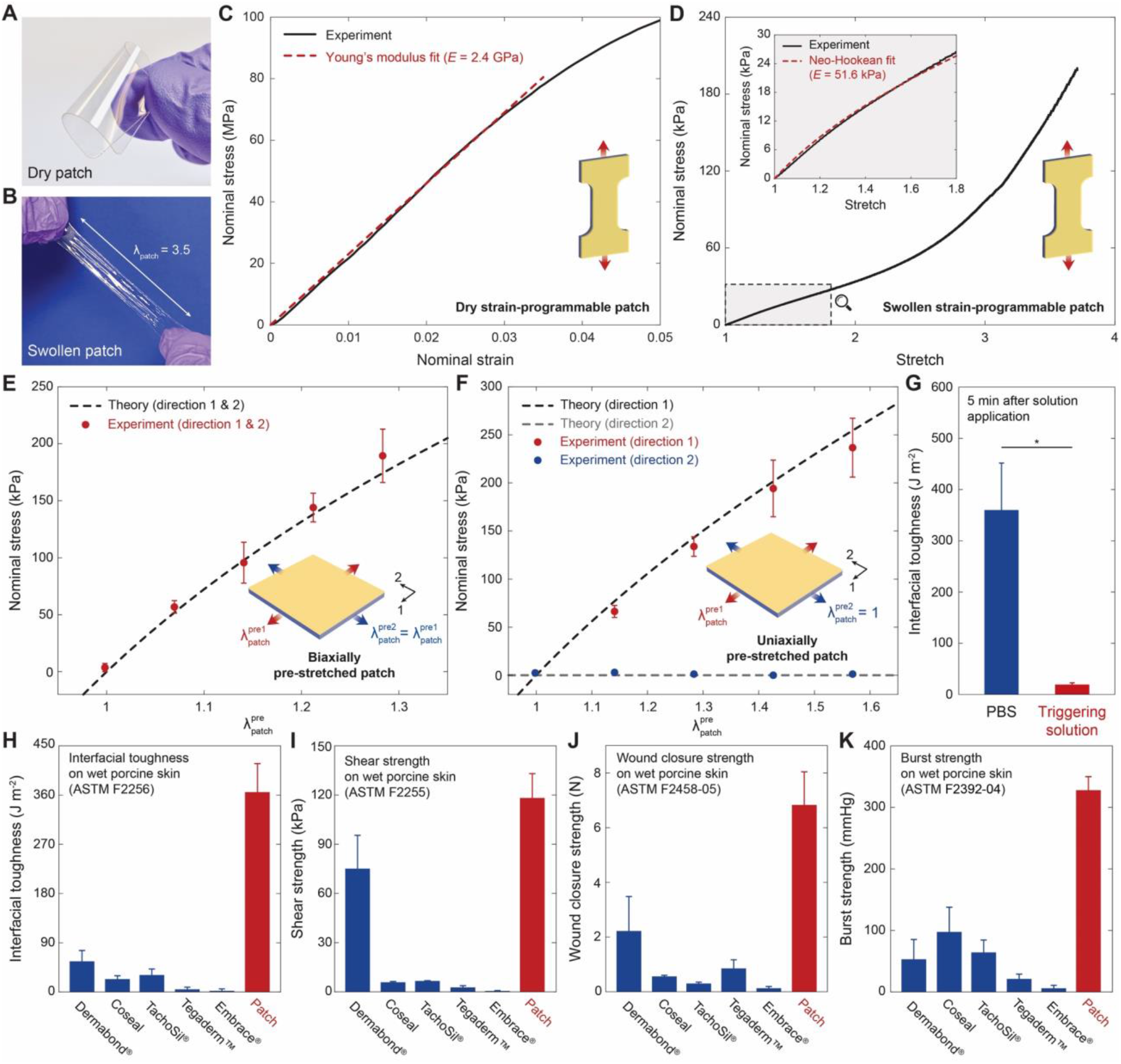
Mechanical properties and adhesion performance. (**A** and **B**) Images of a dry (A) and a swollen (B) strain-programmable patch. (**C** and **D**) Nominal stress vs. stretch curves for a dry (C) and a swollen (D) strain-programmable patch. (**E** and **F**) Experiment and theory values for the nominal contractile stress vs. 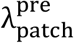 generated by the isotropically (E) and the anisotropically (F) strain-programmed patches. (**G**) Interfacial toughness of the strain-programmable patch on wet porcine skin 5 min after applying PBS and the triggering solution. (**H** to **K**) Interfacial toughness (measured by ASTM F2256) (H), shear strength (measured by ASTM F2255) (I), wound closure strength (measured by ASTM F2458-05) (J), and burst strength (measured by F2392-04) (K) of the strain-programmable patch on wet porcine skin. Values in G to K represent the means ± SD (*n* = 4). *P* values are determined by a Student’s *t* test; ns, not significant; * *p* ≤ 0.05.

Furthermore, the robust interfacial integration between the non-adhesive elastomer backing and the bioadhesive layer in the swollen strain-programmable patch (interfacial toughness over 650 J m^-2^) provides mechanical stability and integrity in wet physiological environments (*30, 31*) (fig. S9). Notably, based on the multi-step pre-stretching fabrication process, the swelling ratio mismatch between the elastomer backing and the bioadhesive layer along the in-plane directions is canceled to minimize the geometric change and the interfacial delamination (figs. S3 and S4; see Supplementary Text for details on the fabrication and strain-programming process).

### Rapid, robust, and on-demand detachable adhesion

To evaluate the adhesion performance of the strain-programmable patch on wet wounded tissues, we conduct four different types of mechanical tests following the testing standards for tissue adhesives to measure the interfacial toughness (by 180-degree peel test, ASTM F2256), shear strength (by lap-shear test, ASTM F2255), wound closure strength (by ASTM F2392-04), and burst strength (by ASTM F2392-04) based on wet porcine skin as the model tissue (*21, 23*) (fig. S10). The strain-programmable patch (with 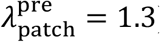) can establish robust adhesion rapidly upon contact and gentle pressure (1 kPa) application for less than 5 s with high interfacial toughness over 350 J m^-2^ (Fig. 2H), shear strength over 115 kPa (Fig. 2I), wound closure strength over 7 N (Fig. 2J), and burst strength over 310 mmHg (Fig. 2K) on wet tissues.

The adhesion performance of the strain-programmable patch outperforms that of commercially-available tissue adhesives and wound dressings including cyanoacrylate adhesives (e.g., Dermabond^®^), polyethylene glycol-based adhesives (e.g., Coseal), fibrin-based adhesives (e.g., TachoSil^®^), elastic wound dressings (e.g., Tegaderm^™^), and contractile wound dressings (e.g., Embrace^®^) (Fig. 2, H to K). Furthermore, taking advantage of the triggerable detachment of the bioadhesive layer (*32*), the adhered strain-programmable patch can be atraumatically detached from the tissue on-demand by applying a biocompatible triggering solution (Fig. 2G, fig. S11, and movie S3). Such benign on-demand removal of the strain-programmable patch can potentially beneficial for the care of chronic diabetic wounds in clinical settings where frequent changes of wound dressings are often required (*33, 34*).

### *In vitro* and *in vivo* biocompatibility

To determine the biocompatibility of the strain-programmable patch and its on-demand detachment process, we perform an *in vitro* cell viability assay based on mouse embryonic fibroblasts (mEFs) and an *in vivo* dorsal subcutaneous implantation based on a rat model (Fig. 3). The *in vitro* biocompatibility of the strain-programmable patch is comparable to that of the control media (DMEM), showing no statistically significant difference in *in vitro* cell viability for mEFs after 24-hour culture (Fig. 3A). The histological assessment made by a blinded pathologist indicates that the strain-programmable patch generates a mild to moderate inflammatory reaction, comparable to or lower than that generated by U.S. Food and Drug Administration (FDA)-approved commercially-available tissue adhesives Coseal and Dermabond^®^ (CA), respectively at 2 weeks after the implantation (Fig. 3, B to D and G). Furthermore, the triggerable detachment process of the strain-programmable patch generates a mild inflammatory reaction comparable to that generated by the sham control group (surgery without implantation) at 2 weeks post-surgery (Fig. 4, E to G).

**Fig. 3.**
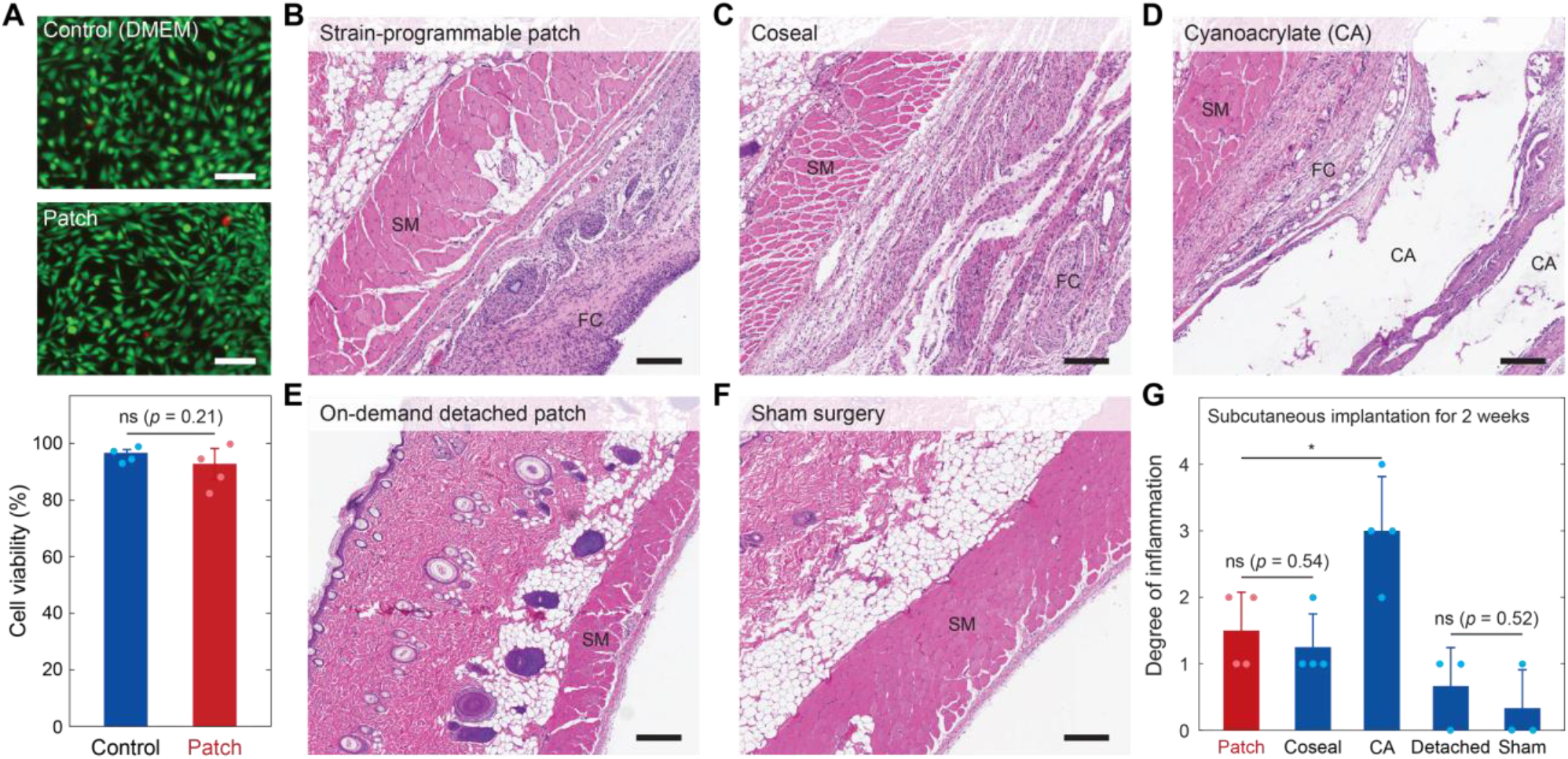
*In vitro* and *in vivo* biocompatibility. (**A**) Representative LIVE/DEAD assay images and the cell viability of mouse embryonic fibroblasts (mEFs) for control (DMEM) and the strain-programmable patch after 24-hour culture. (**B** to **F**) Representative histological images for the subcutaneously implanted strain-programmable patch (B), Coseal (C), Dermabond^®^ cyanoacrylate (CA) adhesive (D), strain-programmable patch after on-demand detachment (E), and sham surgery (F) for 2 weeks stained with hematoxylin and eosin (H&E). (**G**) Degree of inflammation of various groups evaluated by a blinded pathologist (0, normal; 1, mild; 2, moderate; 3, severe; 4, very severe) after 2 weeks of subcutaneous implantation. SM and FC indicate skeletal muscle and fibrous capsule, respectively. All experiments are repeated four times with similar results. Values in A and G represent the means ± SD (*n* = 4). *P* values are determined by Student’s *t* test; ns, not significant; *p* ≤ 0.05. Scale bars, 200 µm (A to F).

**Fig. 4.**
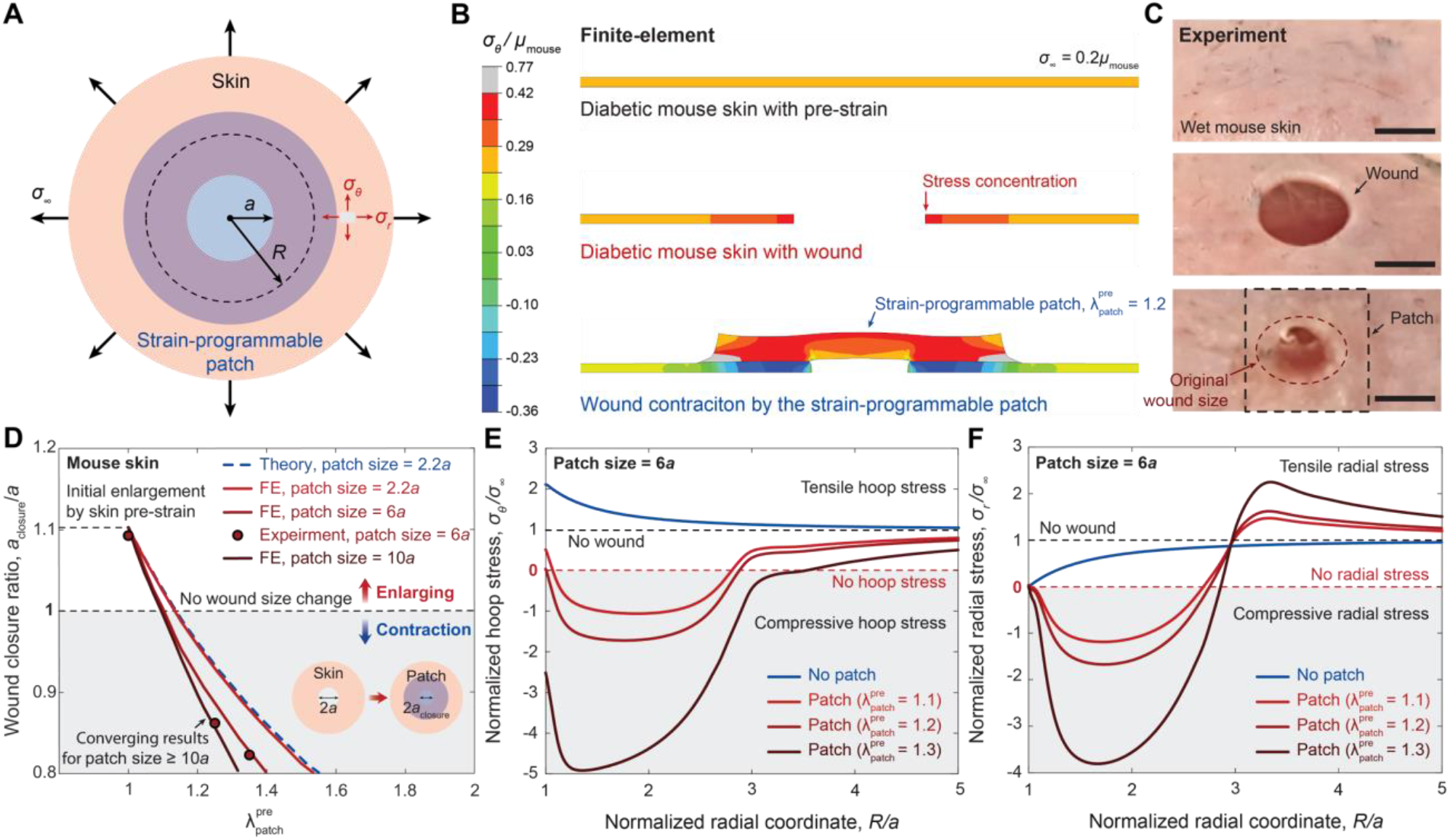
Programmable closure and stress remodeling of diabetic mouse skin wounds. (**A**) Schematic illustration for the theoretical and finite-element analyses. (**B** and **C**) Representative finite-element results (B) and the corresponding experiment images of the *ex vivo* diabetic mouse skin (C) with 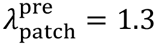 and patch diameter 3 times of the wound diameter. The shear modulus of the diabetic mice skin is denoted as *µ*_mouse_, the hoop stress in the diabetic mouse skin as *σ*_θ_, and the residual stress in the intact diabetic mouse skin as *σ*_∞_. (**D**) Finite-element and experimental results for the wound closure ratio as a function of 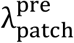 by varying the diameter of the strain-programmable patch. FE, finite-element. (**E** and **F**) Finite-element results for the hoop *σ*_θ_ (E) and the radial *σ*_r_ (F) stresses around the wound with the strain-programmed patches with varying 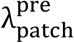. Scale bars, 5 mm (C).

### Programmable mechanical modulation of wound

To provide quantitative guidelines for mechanical modulation of wounds by the strain-programmable patch, we utilize both analytical solutions (figs. S12 and S13) and finite-element method (Fig. 4, A and B, and fig. S14) to model the wound contraction and the remodeling of stresses in the skin around the wound by the strain-programmable patch (see Supplementary Text for details on the analytical and finite-element modeling). Without loss of generality, we study a circular wound in the skin, which is a common form of diabetic wounds. In our models, we take into account a natively existing pre-strain and tension in the skin to better elucidate the mechanical modulation of wounds by the strain-programmable patch (*35-37*). Due to the native pre-strain and tension in the skin, the circular wound undergoes an initial enlargement in the diameter and a substantial increase in hoop stress (over 2 times of the native state) (Fig. 4D and fig. S13, B and D), yielding a stress concentration around the wound edge which can impair wound closure and healing especially in diabetic wounds (*15-17*) (Fig. 4E and fig. S13, C and E). We further validate the model with experiments on *ex vivo* diabetic mouse skin (Fig. 4, A to D) and *ex vivo* human skin (fig. S15, A to D) where the wound closure ratio predicted by the finite-element model shows close agreement with the experimental measurements.

The strain-programmable patch mechanically modulates the wound in diabetic mouse and human skin by (i) reducing the wound diameter (Fig. 4, B to D, fig. S15, B to D, and movie S4) and (ii) reducing the hoop stress around the wound edge (Fig. 4, B and E, fig. S15, C and E) to various degrees based on the relative size of the patch to the wound (Fig. 4D) and the amount of programmed biaxial pre-stretch 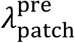. In particular, the strain-programmable patch can effectively reduce the hoop stress concentration around the wound edge at the lower pre-stretch amounts (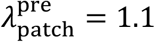 and 1.2), and turn the hoop stress into compressive at the higher pre-stretch amount (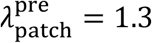) (Fig. 4E and fig. S15E).

### Application on a diabetic mouse model of impaired wound healing

From the above results, we hypothesize that the strain-programmable patch can potentially promote wound healing in diabetic mice at different time frames. In acute or short-term period, the strain-programmable patch can promote diabetic wound healing by applying mechanical contraction and subsequently approximating the wound edges right after the application on wet wounded tissues (Fig. 4, C and D, and movie S4). In chronic or long-term periods, the strain-programmable patch can promote diabetic wound healing by providing a favorable mechanical environment through the remodeling of the stress state around the wound including the reduction in the hoop stress concentration (Fig. 4, B, E, and F).

To assess the efficiency of the patch *in vivo*, we employed an established model of impaired diabetic wound healing; the db/db mouse (*38, 39*). Application of the strain-programmable patch onto 6 mm dorsal excisional wounds resulted in markedly improved wound closure at both 5 and 10 days post-injury (D5 and D10), as evaluated by the % of open wound, degree of re-epithelialization and area of the migrating hyperproliferative neo-epidermis (Fig. 5, A to J) compared to the no strain patch and TD conditions. The strain patch wounds on D10 also displayed well-formed granulation tissue with thick collagen bundles and increased cellularity, which are indicative of advanced healing (Fig. 5D). In addition, strain patch treated wounds exhibited enhanced vascularization as evidenced by the higher density of CD31+ vessels (Fig. 6, A to D). There were also fewer active Caspase-3+ apoptotic cells in the wound bed of strain patch treated mice on D5 (fig. S16C) and more proliferating Ki67+ epidermal cells on D10 wounds (fig. S16E). Consistent with augmented blood vessel formation, extracellular matrix (ECM) production and rapid keratinocyte migration, the gene expression levels of several key pro-angiogenic and pro-healing growth factors were elevated in the strain patch wounds, especially on D5, including *Col3a1, Tgfb1, Vegfa, Fgf2* and *Hgf* (fig. S17A, B, D, and E). Db/db mice have a well-documented deficiency in wound contraction capability (*40*) with dysregulated fibroblast to myofibroblast conversion throughout the healing duration (*41*), so we examined αSMA expression to determine if modifying the wound stress levels via the patch application influences myofibroblast levels (*42*). We found that the strain-programmable patch modified the presence of αSMA+ cells in the wounds, with significantly reduced numbers on D5, but increased on D10 (Fig. 6, E to H). This was accompanied by diminished expression of *Engrailed-1* (*En1*) (fig. S17C and F), a transcription factor recently shown to define an important in wound repair dermal fibroblast subpopulation responsible for fibrosis (*43-45*). We hypothesize that initial strain-programmable patch application stress-shields the tissue and leads to decreased myofibroblasts, but as healing rapidly progresses newly deposited granulation tissue alters the mechanical properties of the wound and activates more αSMA+ cells on D10. Furthermore, adipocyte to myofibroblast differentiation (*46*) or the reverse (*47*) could also be implicated in the higher D10 levels observed as previously reported.

**Fig. 5.**
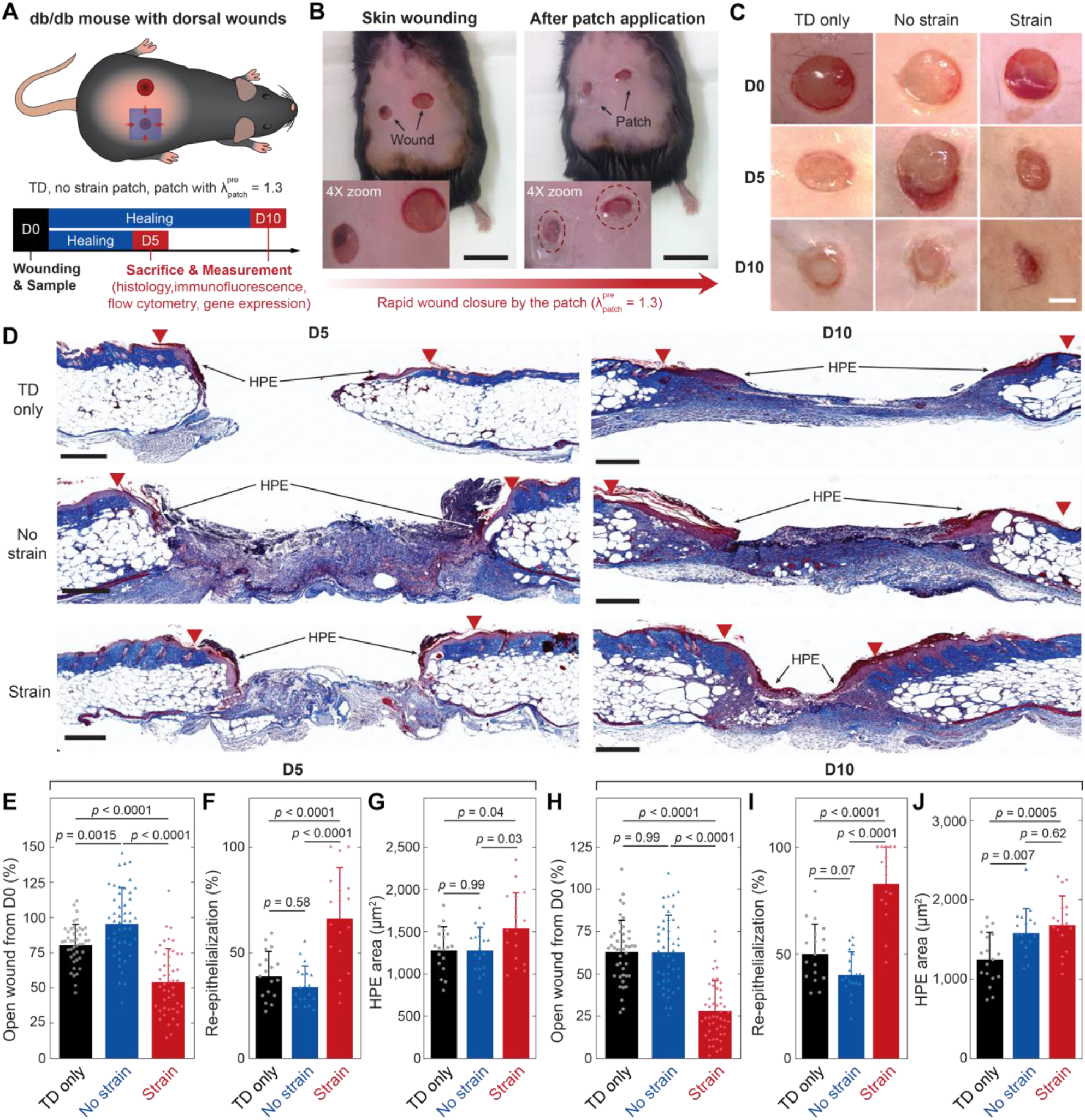
Application of the strain-programmed patch in a mouse model of diabetic wound healing. (**A**) Schematic illustrations for the study design. (**B**) Representative images of db/db mouse dorsal wounds before (left) and after (right) the strain-programmed patch application. (**C**) Representative macroscopic views of wounds on day 0 (D0), day 5 (D5), and day 10 (D10) for Tegaderm™ (TD) only, no strain (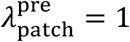) and strain-programmed (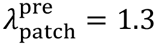) patch experimental groups. The adhered patches were removed before imaging and measurement. (**D**) Representative images from D5 and D10 wounds with Masson’s trichrome stain (MTS). Red triangles denote wound margins; HPE: hyperproliferative epidermis. (**E-G**) Quantification of the wound closure expressed as % of open wound compared to D0 (E), the re-epithelialization expressed as % with 100% being fully covered (F), and the HPE area (G) on D5. (**H-J**) Quantification of the wound closure expressed as % of open wound compared to D0 (E), the re-epithelialization expressed as % (F), and the HPE area (G) on D10. Values in E-J represent the means ± SD (*n* = 16-48 wounds). *P* values were derived from one way ANOVA with Tukey’s post-hoc tests. Scale bars, 20 mm (B); 3 mm (C); 500 µm for D5 and 400 µm for D10 (D).

**Fig. 6.**
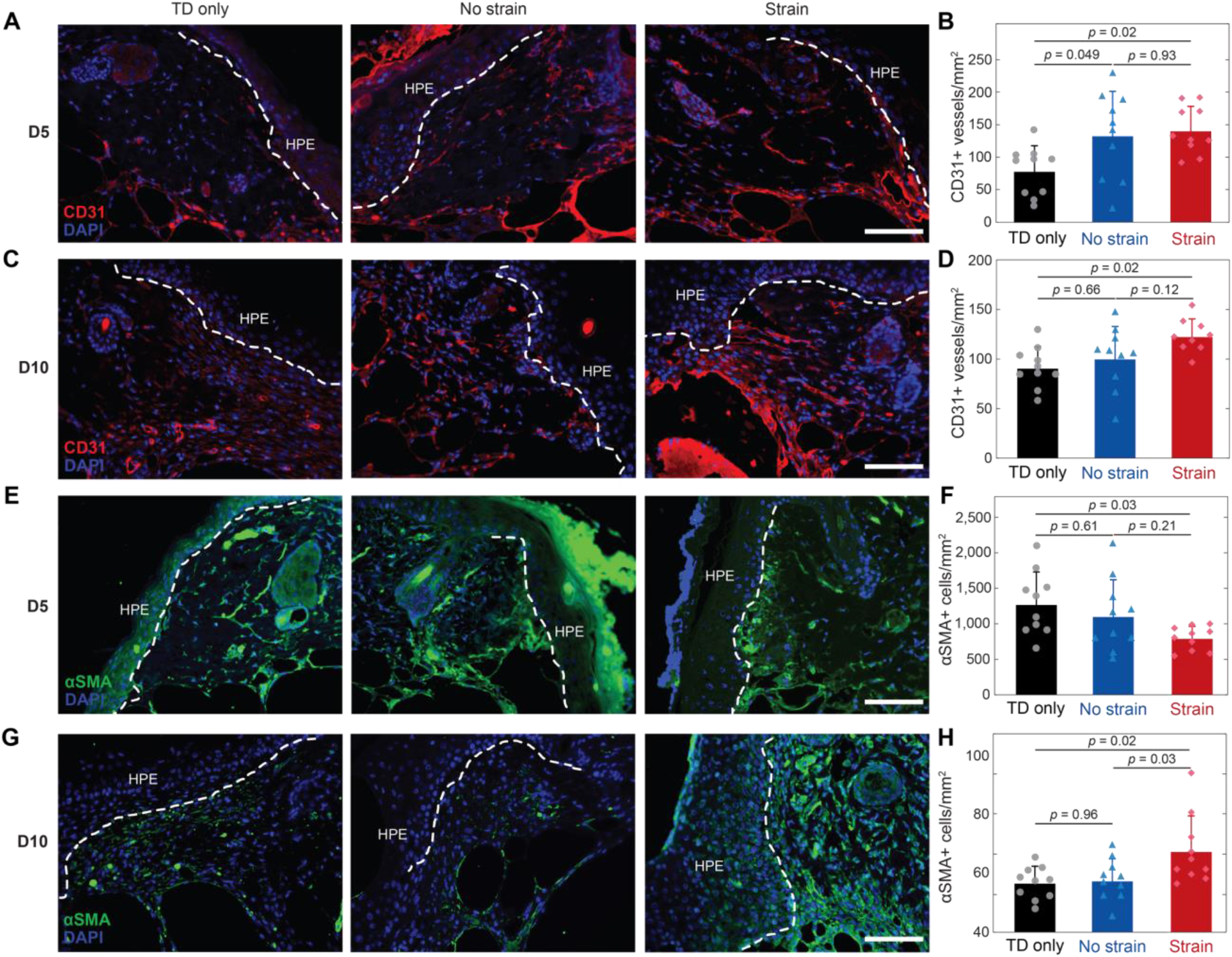
Immunofluorescence staining analysis of diabetic mouse wounds. (**A** and **B**) Representative immunofluorescence images for CD31 (A) and quantification of CD31+ vessels per unit area (B) on day 5 (D5). (**C** and **D**) Representative immunofluorescence images for CD31 (C) and quantification of CD31+ vessels per unit area (D) on day 10 (D10). (**E** and **F**) Representative immunofluorescence images for *α*SMA (E) and quantification of *α*SMA + cells per unit area (F) on D5. (**G** and **H**) Representative immunofluorescence images for *α*SMA (G) and quantification of*α*SMA + cells per unit area (H) on D10. HPE: hyperproliferative epidermis. In immunofluorescence images, blue fluorescence corresponds to cell nuclei stained with 4′,6-diamidino-2-phenylindole (DAPI); red fluorescence corresponds to the expression on blood vessels (CD31); green fluorescence corresponds to the expression on fibroblast (αSMA); dotted line represents the epidermis border. Experimental groups are Tegaderm^™^ (TD) only, no strain (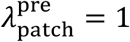) and strain-programmed (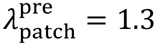) patch for both D5 and D10. Values in B, D, F, H represent the means ± SD (*n* = 9-10 wounds). *P* values were derived from one way ANOVA with Tukey’s post-hoc tests. Scale bars, 100 µm (A); 100 µm (C); 100 µm (E); 100 µm (G).

We then characterized the wound inflammatory cell infiltrate with multicolor flow cytometry (*48*) to profile the major immune cell types affected by the patch application. Our gating strategy is illustrated on figs. S18 and S19. In contrast with the strain and TD groups, there were more total immune cells (CD45+) in no strain patch treated wounds on D5 (fig. S20 A), including more neutrophils (CD45+CD64-Ly6G+) (fig. S20B) and monocytes (CD45+CD11b+CD64-/intLy6C+) (fig. S20C), but fewer macrophages (CD45+CD11b+CD64+F4/80+) (fig. S20D) and T-cells (CD45+CD3+) (fig. S20F). The strain patch also induced an amplified immune response but to a moderate extent compared with the no strain patch. This is to be expected, as any interaction of a biomaterial with the immune system triggers an immune response (*49*). We also analyzed the expression of established macrophage polarization markers and discovered that on D5 the patch treated groups displayed an M1-skewed phenotype, with increased % of CD80 and CD86 M1 macrophages (fig. S20I and J) and reduced % of CD163 and CD301b M2 macrophages (fig. S20K and M). Interestingly, they also had higher % of typically M2 associated CD206 cells (fig. S20L). On D10, there were more immune cells and neutrophils in the strain and no strain wounds (fig. S21A and B) and more monocytes and fewer macrophages in the no strain wounds (fig. S21C and D). The sustained increased number of monocytes and reduced macrophages denotes insufficient monocyte to macrophage differentiation and a prolonged inflammatory state. In addition, macrophages in the strain patch-treated wounds had a more M2-like phenotype with decreasing % of CD80 (fig. S21J) and increasing % of CD163 and CD301b (fig. S21K and M) suggesting that the healing was actively progressing towards the proliferation phase (*50*).

Next, to better understand the mechanisms of the observed wound healing acceleration, we performed bulk RNA-seq on D10 wound tissues. Principal component analysis (PCA) revealed separate clusters of the samples according to treatment, indicating distinct transcriptome profiles (Fig. 7A). Differential gene expression analysis with log(fold change) < 1 or > 1 and false discovery rate (FDR) < 0.05 on strain-programmable patch treated wounds versus Tegaderm only identified 3,581 significantly modified genes (1681 upregulated) (Fig. 7B) and 62 genes (14 upregulated) in strain-programmable versus no strain patch (Fig. 7C). The volcano plots and heat maps (fig. S22) illustrate the most highly expressed features, while the complete lists are presented in Data file S1. Further, overrepresentation analysis of the top differentially expressed genes highlighted the enrichment of multiple processes linked to muscle contraction which is in agreement with our observation of more αSMA+ cells in the strain-programmable patch wounds by D10 (Fig. 7, D to G). A similar analysis for no strain versus TD yielded cytokine related pathways as most enriched (fig. S23). Collectively, these findings indicate that applying the patch on diabetic murine cutaneous wounds promotes healing by positively affecting multiple integral reparative processes, including keratinocyte migration, angiogenesis and proliferation. It also alleviates the tension of the tissue leading to an initial diminished and subsequently increased myofibroblast presence which also proves beneficial for wound closure.

**Fig. 7.**
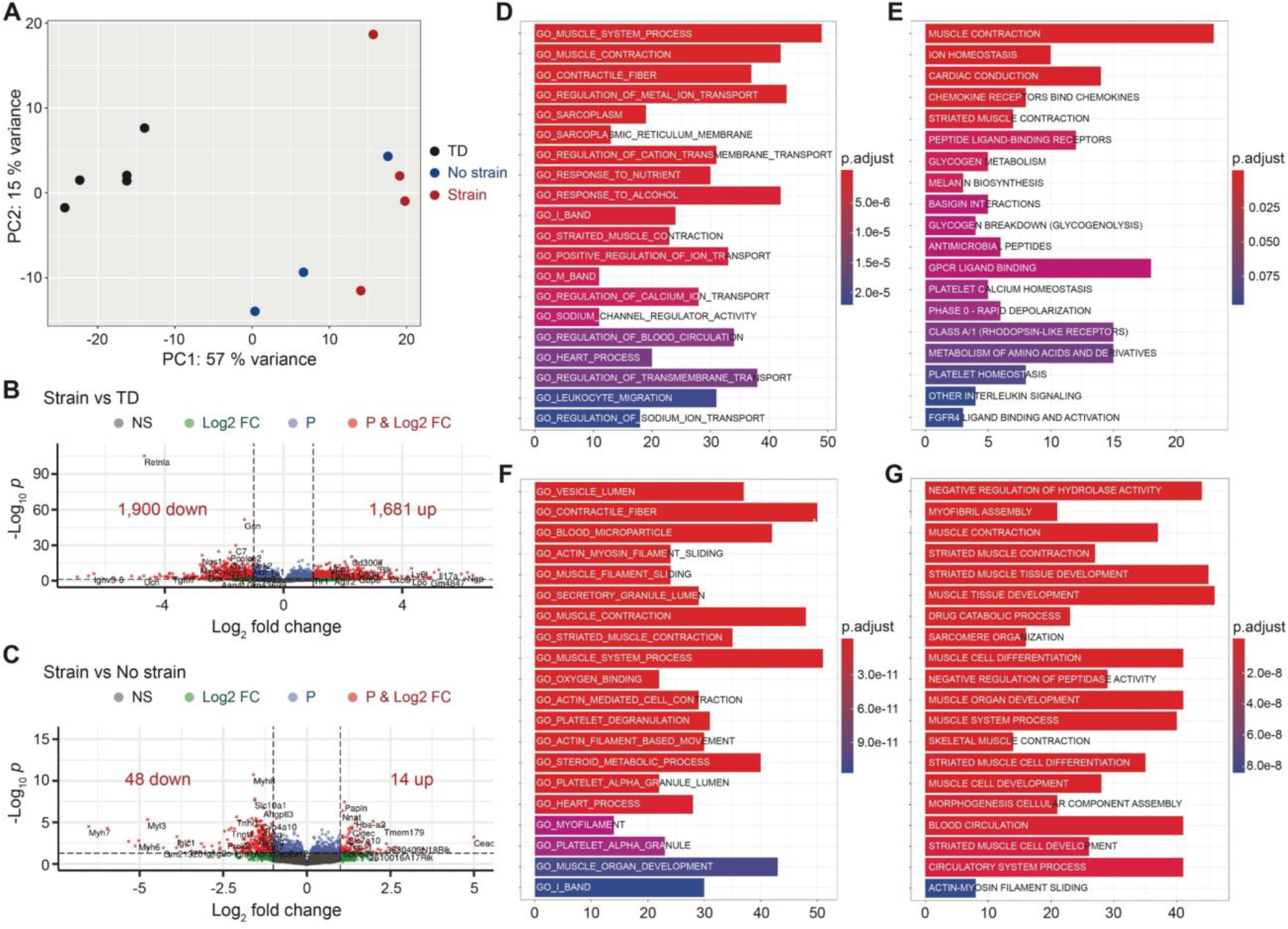
Significantly altered global transcriptome of db/db mouse skin wounds. (**A**) Principal component analysis (PCA) plot illustrating the variances of the Tegaderm™ (TD) only (black dots, *n* = 5), no strain (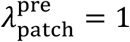) (blue dots, *n* = 3) and strain-programmed (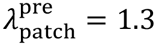) (red dots, *n* = 4) patch datasets. (**B** and **C**) Volcano plots displaying gene expression profiles when comparing the strain-programmed patch against TD (B) and the strain-programmed against no strain (C) patches. Red colored data points represent genes that meet the thresholds of fold change (FC) above 1 or under −1, False Discovery Rate (FDR) < 0.05. (**D** to **G**) Functional over-representation analysis utilizing the top 500 differentially expressed genes results for Strain vs TD and Strain vs No strain in gene ontology (GO) (D and F) and Reactome (E and G) databases. The x-axis corresponds to the number of genes implicated in each pathway and the color of the bars correlates with the adjusted *p*-values as shown in the legends.

### Application on a human *ex vivo* model of wound healing

To examine potential effects of the patch on human wound healing, we inflicted 6 mm punch biopsy wounds on panniculectomy derived discarded skin kept in cell culture conditions (*51, 52*) and monitored healing over four days (Fig. 8A and B). We quantified the distance between the two edges of the migrating epidermis as a measure of wound healing and found the strain patch to promote faster re-epithelialization compared to the other two conditions (Fig. 8C and E). Masson’s trichrome staining (MTS) for the assessment of collagen fibers and scoring by an experienced pathologist demonstrated elevated intensity in the strain patch-treated wounds suggesting that the strain-programmable patch also influences the dermal ECM (Fig. 8D and F). We observed no differences in the number of fibroblast-like cells or vessels counted from hematoxylin and eosin (H&E) stained wound sections (Fig. 8G and H).

**Fig. 8.**
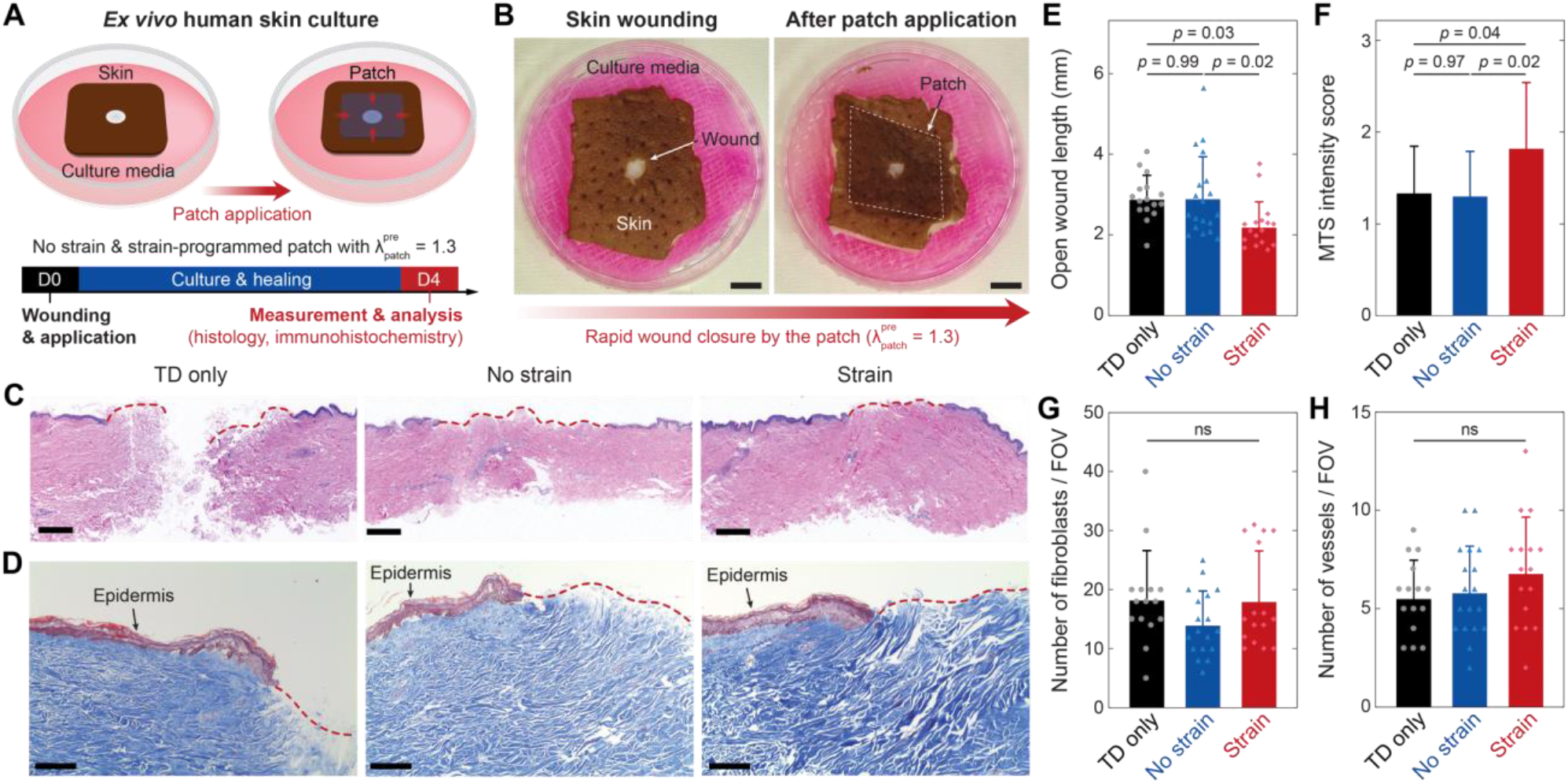
Application of the strain-programmed patch in a human *ex vivo* model of wound healing. (**A**) Schematic illustrations for the study design. (**B**) Representative images of an *ex vivo* human skin culture setup before (left) and after (right) the strain-programmed patch application. (**C** and **D**) Representative images from day 4 (D4) wounds for Tegaderm^™^ (TD) only, no strain (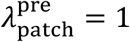) and strain-programmed (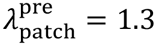) patch experimental groups with hematoxylin and eosin stain (H&E) (C) and Masson’s trichrome stain (MTS) (D). Red dotted line marks the non-epithelized tissue. (**E-H**) Quantification of the open wound length (E), the MTS intensity score (F), the number of fibroblast-like cells per field of view (FOV) (G), and the number of vessels per FOV (H) on D4. Values in E-H represent the means ± SD (*n* = 15-19 wounds from 3 individual patients’ skin). *P* values were derived from one way ANOVA with Tukey’s post-hoc tests. Scale bars, 10 mm (B); 800 µm (C); 200 µm (D).

To further explore the efficiency of the patch clinically and taking into consideration that DFU can be irregularly shaped, we utilized two representative examples of images taken from patients at the Joslin-Beth Israel Deaconess Foot Center and recreated similar wounds on *ex vivo* porcine skin (fig. S25A and B). Digital photography and FE simulation following the strain-programmable patch application revealed that initial wound area was decreased and tension relief was achieved (fig. S25 C to F), which are encouraging insights into its future use in the clinic for treating DFU.

## DISCUSSION

Our results show that application of the strain-programmable patch on db/db mouse wounds achieves a 50% wound closure by day 5 post-injury and a remarkable 75% on day 10 and thus outperforms the previously reported interventions in similar models (*53-55*), including PDGF treatment (*56*) which is the only recombinant growth factor FDA-approved DFU therapy (*57*). Our findings that modulation of wound tension ameliorates healing complement and expand on the recent work by Mascharak *et al*. (*44*). They used both a device to control wound mechanical forces and a pharmacological inhibition strategy to demonstrate that *Engrailed-1* expressing fibroblasts activated via mechanotransduction signaling are responsible for fibrosis in the wound and blocking them results in regeneration. Here, we show that a similar approach is also highly effective in impaired diabetic wound healing.

Profiling the major immune cell types involved in wound healing we discover that the strain-programmable patch elicits an inflammatory response compared to the Tegaderm control, which however does not prove detrimental to wound closure. Chronic diabetic wounds, like the DFU, are mainly characterized by the persistence of low-grade inflammation and inability to progress to the next phase of wound healing (*18, 58-60*). Previous studies in our unit have shown that approaches that improve wound healing in diabetic murine models exert their beneficial effects by converting the low-grade chronic inflammation to an acute one and that this conversion is adequate to promote linear progression to the next phases and accelerate wound healing (*61, 62*). Similar findings have also been reported by us and others in human studies (*63, 64*).

FDA has approved four products for DFU treatment and all of them were developed in the 1990s: becaplermin (rhPDGF-BB), a recombinant growth factor (*6*); two bioengineered skin substitutes (Apligraf (*4*) and Dermagraft (*5*)); and Omnigraft that is based on Integra Dermal Regeneration Matrix (*65*). However, the efficacy of these products is rather limited, as in the pivotal trials, half or more of the participants failed to heal their wounds (*66, 67*) while their cost is considerable. Numerous other clinical trials with additional growth factors, including bFGF, EGF, and VEGF, devices, and other techniques have all failed to show meaningful efficacy (*68*). Furthermore, although basic research studies have indicated that DPP4 inhibitors may promote wound healing and reduce fibrosis (*43*), there is no concrete clinical evidence that these inhibitors can have any effect of DFU healing. This underscores the significant need to develop new therapies that could be tested in future clinical trials.

It has been well-established that mechanical reinforcement or stimuli to alleviate the adverse stress concentration around the wound can facilitate wound healing of healthy skin both in animal models and human clinical trials (*10-12, 17, 60, 69*). However, any potential therapeutic effects of such mechanical stimuli are underexplored in chronic diabetic wounds such as DFU with impaired wound healing including a reduced degree in contractility of wound edges and subsequent closure of wounds (*18, 58-60, 66*). We hypothesize that a tissue adhesive biomaterial capable of providing programmed mechanical contractions can facilitate the healing of diabetic wounds by addressing this mechanical imbalance. To implement this hypothesis into a viable therapeutic platform, we develop the strain-programmable patch that synergistically incorporates the hydration-based shape-memory mechanism and the dry-crosslinking mechanism to achieve rapid, robust, and fully programmable mechanical modulation of wet wounded tissue. We provide a quantitative design guideline for the predictable and rational design of the proposed strain-programmable patch based on theoretical, numerical, and experimental analysis and modeling. We further validate biocompatibility and diabetic wound healing efficacy of the strain-programmable patch via *in vitro* and *in vivo* rodent models. Taking advantage of the quantitative design guideline and fabrication process for the strain-programmable patch, we also demonstrate that the strain-programmable patch can be readily translated into human-scale diabetic wound healing applications based on an *ex vivo* human skin wound healing model.

Overall, our *in vivo* rodent and *ex vivo* human diabetic wound healing data validate our hypothesis and suggest that the strain-programmable patch can offer a promising therapeutic solution for the treatment of chronic diabetic wounds. We envision that the strain-programmable patch has the potential for commercialization and eventual translation as a clinical treatment for chronic diabetic wounds. While the current work provides systematically investigated platform technology and proof-of-concept efficacy of the therapeutic potential, there are future steps to be taken for further investigation and clinical translation. Future investigations should focus on employing multi-omics methods to comprehensively map the different cell populations and signaling pathways influenced by the strain programmable-patch treatment (*70, 71*). In addition, further studies are required to define applicability, frequency of application and suitable strain levels, among others, before use in a clinical setting. Completion of this work has the potential to lead to phase I/II clinical trials.

## MATERIALS AND METHODS

### Study design

The aim of this study was to develop a strain-programmable bioadhesive patch to provide an effective, reliable, and precisely-controllable mechanical modulation of wet diabetic wounds for accelerated healing. It was hypothesized that a bioadhesive material capable of both rapid robust adhesion on wet wounded tissues and programmable contraction would facilitate the healing of diabetic wounds by addressing their inherent mechanical imbalance. Mechanical characterizations were performed based on *ex vivo* porcine skin to optimize and evaluate the adhesion performance and strain-programmability of the proposed platform. *In vitro* tests were performed to evaluate the cytotoxicity of the strain-programmable patch. *In vivo* biocompatibility was evaluated by a blinded pathologist based on a rat subcutaneous implantation model. Diabetic wound healing efficacy of the strain-programmable patch was evaluated *in vivo* based on the db/db mice wound healing model. Clinical translational potential was assessed with an *ex vivo* human skin culture model. The efficacy of diabetic wound healing in comparison with a standard care control group (Tegaderm™, 3M) was evaluated based on histology, immunohistochemistry, flow cytometry, RNA-seq and gene expression analyses.

### Preparation of the strain-programmable patch

To prepare the bioadhesive layer, acrylic acid (30 w/w %), chitosan (HMC+ Chitoscience Chitosan 95/500, 95 % deacetylation, 2 w/w %), α-ketoglutaric acid (0.2 w/w %), and poly(ethylene glycol methacrylate) (PEGDMA; Mn = 550, 0.03 w/w %) were dissolved in deionized water. Then, we dissolved 100 mg functional monomer (NHS ester functionalized monomer with disulfide bond) synthesized following the previous report (*32*) in 1 ml acetone and added to 10 ml of the above stock solution to get a precursor solution. The precursor solution was then poured on a glass mold with spacers (the thickness is 210 μm unless otherwise mentioned) and cured in ultraviolet light chamber (365 nm, 10 W power) for 30 min. As a non-adhesive elastomer backing resin, 10 w/w % hydrophilic polyurethane (AdvanSource Biomaterials) in ethanol/water mixture (95:5 v/v) was spin-coated on the as-prepared bioadhesive at various speed (fig. S25, 100 rpm was used otherwise mentioned).

The as-prepared bioadhesive layer coated with the non-adhesive elastomer backing resin underwent multi-step processes to fabricate the strain-programmable patch (fig. S3). Detailed fabrication steps were described in the Supplementary Information. The prepared dry strain-programmable patch was sealed in a plastic bag with desiccants (silica gel packets with active charcoal, McMaster Carr) and stored at –20 °C before use. To prepare the triggering solution, 0.5 M sodium bicarbonate (SBC) and L-glutathione reduced (GSH) were dissolved in phosphate buffered saline (PBS). The triggering solution was filtered by using a 0.2-µm sterile syringe filter before use.

### Mechanical characterization

Unless otherwise indicated, the strain-programmable patch (with 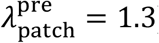) was applied after washout of the tissue surfaces with PBS followed by 5 s pressing (with 1 kPa pressure applied by either mechanical testing machine or equivalent weight). Unless otherwise indicated, all mechanical tests on adhesion samples were performed 12 h after initial pressing to ensure equilibrium swelling of the adhered strain-programmable patch in wet environments. The application of commercially-available tissue adhesives and wound dressings followed the provided manual for each product.

To measure interfacial toughness, adhered samples with widths of 2.5 cm were prepared and tested by the standard 180-degree peel test (ASTM F2256) using a mechanical testing machine (2.5-kN load-cell, Zwick/Roell Z2.5). All tests were conducted with a constant peeling speed of 50 mm min^-1^. The measured force reached a plateau as the peeling process entered the steady-state. Interfacial toughness was determined by dividing two times the plateau force by the width of the tissue sample (fig. S10A). Hydrophilic nylon filters (1 µm pore size, TISCH Scientific) were applied as a stiff backing for the strain-programmable patch. Poly(methyl methacrylate) (PMMA) films (with a thickness of 50 µm; Goodfellow) were applied using cyanoacrylate glue (Krazy Glue) as a stiff backing for the tissues. For on-demand detachment of the strain-programmable patch, the interfacial toughness was measured 5 min after applying the triggering solution.

To measure shear strength, the adhered samples with an adhesion area of 2.5 cm in width and 1 cm in length were prepared and tested by the standard lap-shear test (ASTM F2255) with a mechanical testing machine (2.5-kN load-cell, Zwick/Roell Z2.5) (fig. S10B). All tests were conducted with a constant tensile speed of 50 mm min^-1^. Shear strength was determined by dividing the maximum force by the adhesion area. Hydrophilic nylon filters were applied as a stiff backing for the strain-programmable patch. PMMA films were applied using cyanoacrylate glue (Krazy Glue) as a stiff backing for the tissues.

To measure wound closure strength, the adhered samples with 2.5 cm in width and 1 cm in overlap length (between adhesive and tissue) were prepared and tested by the standard wound closure strength test (ASTM F2458-05) with a mechanical testing machine (2.5-kN load-cell, Zwick/Roell Z2.5) (fig. S10C). All tests were conducted with a constant tensile speed of 50 mm min^-1^. Wound closure strength was determined by measuring the maximum force.

To measure burst strength, the sealed samples with 3-mm hole were prepared and tested by the standard burst strength test (ASTM F2392-04) with a pressure gauge (Omega) (fig. S10D). All tests were conducted with a constant flow rate of PBS at 2 mL min^-1^. Burst strength was determined by measuring the maximum pressure.

The tensile properties and fracture toughness of the strain-programmable patch were measured using pure-shear tensile tests of thin rectangular samples (10 mm in length, 30 mm in width, and 0.5 mm in thickness) with a mechanical testing machine (20-N load-cell, Zwick/Roell Z2.5). All tests were conducted with a constant tensile speed of 50 mm min^-1^. The fracture toughness of the strain-programmable patch was calculated by following the previously reported method based on tensile tests of unnotched and notched samples (*72*) (fig. S7).

The tensile properties of *ex vivo* diabetic mouse skin and human skin were measured with a mechanical testing machine (2.5-kN load-cell, Zwick/Roell Z2.5). All tests were conducted with a constant tensile speed of 50 mm min^-1^. The nominal stress vs stretch curves of skin were fitted with the incompressible Ogden hyperelastic model as

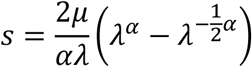

where *s* is nominal stress (i.e., the measured force divided by the cross-sectional area of an undeformed sample), *µ* is shear modulus, and *α* is Ogden coefficient. For diabetic mice skin, *μ*_mouse_ = 87.5 kPa and *α*_mouse_ = 7. For human skin, *μ*_human_ = 40 kPa and *α*_human_ = 20 (fig. S26).

### *In vitro* biocompatibility

*In vitro* biocompatibility tests were conducted by using the strain-programmable patch-conditioned media for cell culture (*73*). To prepare the strain-programmable patch-conditioned media for *in vitro* biocompatibility tests, 20 mg of the strain-programmable patch was incubated in 1 mL Dulbecco’s modified eagle medium (DMEM) at 37 °Cfor 24 h. The pristine DMEM was used as a control. Wild-type mouse embryonic fibroblasts (mEFs) were plated in 96-well plate (*N* = 6 per each case). The cells were then treated with the strain-programmable patch-conditioned media and incubated at 37 °Cfor 24 h in 5 % CO_2_. The cell viability was determined with a LIVE/DEAD viability/cytotoxicity kit for mammalian cells (Thermo Fisher Scientific) by adding 4 µM calcein and ethidium homodimer-1 into the culture media. A confocal microscope (SP 8, Leica) was used to image live cells with excitation/emission at 495nm/515nm, and dead cells at 495nm/635nm, respectively. The cell viability was calculated by counting live (green fluorescence) and dead (red fluorescence) cells by using ImageJ.

### *Ex vivo* skin study

All *ex vivo* experiments were reviewed and approved by the Committee on Animal Care at the Massachusetts Institute of Technology. For closure of isotropic porcine skin wounds, a hole was made on a porcine belly with an 8-mm biopsy punch. For closure of anisotropic porcine skin wounds, a 3-cm long laceration was made on a porcine belly with a scalpel. For closure of diabetic mouse skin wounds, a hole was made on a dorsal skin with a 6-mm biopsy punch. The strain-programmable patch with 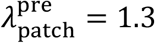 (for both biaxial and uniaxial pre-stretch) was used for *ex vivo* study.

### *In vivo* biocompatibility

All animal surgeries were reviewed and approved by the Committee on Animal Care at the Massachusetts Institute of Technology. Female Sprague Dawley rats (225-250 g, Charles River Laboratories) were used for all *in vivo* studies. Before implantation, the strain-programmable patch was prepared using aseptic techniques and was further sterilized for 3 h under UV light. For implantation in the dorsal subcutaneous space, rats were anesthetized using isoflurane (1–2% isoflurane in oxygen) in an anesthetizing chamber. Anesthesia was maintained using a nose cone. The back hair was removed and the animals were placed over a heating pad for the duration of the surgery. The subcutaneous space was accessed by a 1-2 cm skin incision per implant in the center of the animal’s back. To create space for implant placement, blunt dissection was performed from the incision towards the animal shoulder blades. For the sham surgery group, no implant was placed in the subcutaneous pocket (*n* = 4). For the triggerable detachment group, the strain-programmable patch (10 mm in width and 20 mm in length) was placed in the subcutaneous pocket created above the incision and detached 5 min after applying 1 mL of the triggering solution (*n* = 4). For the patch group, the strain-programmable patch (10 mm in width and 20 mm in length) was placed in the subcutaneous pocket created above the incision without detachment (*n* = 4). For commercially-available tissue adhesive groups, 0.5 mL of Coseal (*n* = 4) and Dermabond^®^ cyanoacrylate adhesive (*n* = 4) were injected in the subcutaneous pocket created above the incision. The incision was closed using interrupted sutures (4-0 Vicryl, Ethicon) and 3-6 ml of saline were injected subcutaneously. Up to four implants were placed per animal ensuring no overlap between each subcutaneous pocket. After 2 weeks following the implantation, the animals were euthanized by CO_2_ inhalation. Subcutaneous regions of interest were excised and fixed in 10 % formalin for 24 h for histological analyses.

Fixed tissue samples were placed into 70 % ethanol and submitted for histological processing and hematoxylin and eosin (H&E) staining at the Hope Babette Tang (1983) Histology Facility in the Koch Institute for Integrative Cancer Research at the Massachusetts Institute of Technology. Histological assessment was performed by a blinded pathologist on a scale of 0-4 (0, normal; 1, mild; 2, moderate; 3, severe; 4, very severe) to evaluate the degree of inflammation in the tissues surrounding the subcutaneous implants. The degree of acute inflammation was based on the number of neutrophils. The degree of chronic inflammation was based on the presence of lymphocytes, macrophages, and plasma cells. The degree of inflammation was evaluated based on the overall presence of indicators in each histological sample (normal, mild, moderate, severe, very severe). Representative images of each group were shown in the corresponding figures.

### *In vivo* diabetic mice wound healing study

All procedures were approved by the BIDMC Institutional Animal Care and Use Committee. Male db/db mice (stock # 000642) were obtained from Jackson Laboratories and were acclimated to the animal facility for at least one week before surgery. They were routinely weighed and their blood glucose was assessed with a commercially available glucometer (Contour, Bayer) and confirmed to be > 250 mg dL^-1^. 12-week old mice were anesthetized using isoflurane and two circular biopsy punch 6-mm (Integra Miltex) full-thickness wounds were created on their depilated and disinfected dorsum. Sterile strain programmable patches, no strain patches or no patches were immediately placed on the wounds. The wounds were then covered with an occlusive dressing (Tegaderm, 3M) for protection. The mice were housed individually after surgery and observed every day until euthanized with excess CO_2_ on days 5 or 10. The wounds were photographed on days 0, 5 and 10 with a standard iphone5 camera secured on a stand and measured with digital calipers (Thermo Fisher, 14-648-17). A ruler was placed beside the wounds as a scale bar for area calculation. Wound closure was quantified using both ImageJ and calipers measurements and expressed as percentage healed compared to day 0.

### *Ex vivo* human skin culture wound healing study

Skin specimens were obtained from BioIVT (Westbury, NY) and derived from abdominoplasty procedures of three female patients aged 28 to 41. De-identified samples were provided with removed subcutaneous adipose tissue in sterile PBS at 4° C on the day of surgery and were immediately processed. Prior to wounding, skin specimens were sterilized by sequential washes with iodine, 70% ethanol, and PBS and were then cut into evenly sized squares to fit into a 60 mm tissue culture dish (ThermoFisher). A 6 mm biopsy punch was used to gently punch the skin’s epidermis partially penetrating into the dermis to create a wound. Strain-programmable patches, no strain patches or Tegaderm were applied onto the wounds. A sterile gauze was placed on top of the tissue culture dish and soaked with high glucose DMEM (ThermoFisher) supplemented with 10% heat inactivated FBS (Sigma-Aldrich) and 1% Penicillin-Streptomycin (Gibco). Skin specimens were next placed dermis side down onto the culturing dishes and incubated at 37 °C, 5% CO_2_. Media was changed every day.

### Histology immunohistochemistry and immunofluorescence

Human or mouse wound tissues following completion of the study on days 4 or 5 and 10 respectively were bisected at the wound center. One half was snap-frozen and stored at -20 °C, while the other was fixed in 10 % formalin and processed for paraffin embedding. 5 µm thick sections were used. MTS and H&E stains were performed at BIDMC Histology Core. Whole slide image acquisition was performed at the DF/HCC Research Pathology Cores with an Aperio CS2 scanner (Leica Biosystems). Re-epithelialization was quantified from MTS images by measuring the length of the migrating epithelial tongue covering the wound and dividing by the entire length of the wound. An experienced dermatopathologist (A.K.) scored the human wounds MTS slides according to intensity on a scale of 1 to 3 and counted the number of fibroblast-like cells and blood vessels. For immunohistochemistry of mouse wounds, tissue sections were deparaffinized, rehydrated and antigen retrieval was achieved with citrate buffer pH 6.0 in a pressure cooker for 15 min. They were then blocked in 1% BSA for 1.5 h at RT. The Vectastain Elite ABC Rabbit IgG Kit (Vector Laboratories) was used following the manufacturer’s instructions. Sections were incubated with rabbit anti-active caspase-3 (1:40, BD, 559565) overnight at 4 °C. Visualization of the secondary biotinylated antibody binding was performed using NovaRED substrate kit (Vector Laboratories). Images of sections were obtained at 10x magnification with Elipse E200 upright microscope (Nikon) using Motic Images Plus 3.0 software.

For immunofluorescence staining, deparaffinization and antigen retrieval of the paraffin-embedded mice wound sections were performed as previously described. The primary antibodies used were: mouse monoclonal anti-Cytokeratin 14 (1:1000, Abcam ab7800), rat monoclonal anti-Ki67 (1:200, eBioscience, 14-5698-82), rabbit polyclonal anti-CD31 (1:50, Abcam ab28364) and goat polyclonal anti-alpha smooth muscle actin (1:100, Abcam ab21027). Sections were first blocked (5% donkey serum, 1% BSA in 0.1% Triton-X PBS) for 1 hr at RT and then incubated with primary antibodies overnight at 4 °C in a humidified chamber. The secondary antibodies used were all donkey at 1:500 dilution: anti-mouse IgG H&L (Alexa Fluor® 488, ab150109); anti-rat IgG H&L (Alexa Fluor® 647, ab150155), anti-rabbit IgG H&L (Alexa Fluor® 594, ab150064); anti-goat IgG H&L (Alexa Fluor® 488, ab150133); anti-rabbit IgG H&L (Alexa Fluor® 594, ab150064); anti-foat IgG H&L (Alexa Fluor® 647, ab150131). Nuclear counterstaining was performed with DAPI (Invitrogen). Sections were quenched for 5 minutes using the TrueView Autofluorescence Quenching kit to decrease background (Vector Laboratories) and covered with anti-fade mounting medium (Vector Laboratories). Images were obtained at 10x and 20x magnification with an Axio Imager A2 microscope using Zen Blue edition software (Zeiss). Quantification of the stainings was performed on Image J/FIJI by counting the positive cells/structures for a particular marker and dividing by the area of the tissue for normalization.

### Skin dissociation and flow cytometry

Immediately following sacrifice, mouse skin comprising of wound and approximately 0.5 mm of peri-wound tissue was excised and kept on ice cold sterile PBS until processing within 2 hr. Four wounds from two mice were pooled as one sample to ensure enough single cells. The skin was finely minced with a scalpel and placed for 30 min at 37 °C on a shaker in a digesting enzyme cocktail of 2 mg/ml Collagenase P (Roche), 2 mg/ml Dispase (Gibco) and 1 mg/ml DNase I (Stemcell Technologies) in DMEM (Gibco) with 10% FBS and 1% P/S, using glass pipettes to break down the ECM every 10 min. Single cell suspensions were passed through a 40 μm cell strainer, counted with a K2 cellometer (Nexcelom Bioscience) and cryopreserved in 90% FBS 10% DMSO until processing.

Cells were quickly thawed and adjusted to a concentration of 10^6^ cells/mL. A LIVE/DEAD fixable dead cell stain kit was used to exclude dead cells from the analysis (ThermoFisher). AbC total antibody and amine reactive ArC compensation beads kits (ThermoFisher, A10497 and A10346) were included for single stain controls. After completing the viability stain per the manufacturer’s instructions, cells were blocked (Biolegend FACS buffer with 0.05% anti-mouse CD16/32 Biolegend, 101320 and 0.05% Tru-stain monocyte blocker Biolegend, 426103) for 10 min at RT. An antibody cocktail with details listed on Data file S2 was then added for 25 min on ice. Cells were washed 2x, fixed with 0.4% PFA and stored at 4° C protected from light until analysis the next day. The samples were run on a CytoFLEX LX flow cytometer (Beckman Coulter) and data was processed and analyzed with CytExpert software (Beckman Coulter) at the BIDMC Flow Cytometry Core.

### RNA extraction, sequencing and analysis

RNA extraction, library preparations, and sequencing reactions were conducted at GENEWIZ, LLC. (South Plainfield, NJ, USA). Total RNA was extracted using the Qiagen RNeasy Plus Universal mini kit following the manufacturer’s instructions (Qiagen, Hilden, Germany). Extracted RNA samples were quantified using Qubit 2.0 Fluorometer (Life Technologies, Carlsbad, CA, USA) and RNA integrity was checked on Agilent TapeStation 4200 (Agilent Technologies, Palo Alto, CA, USA). RNA sequencing libraries were prepared using the NEBNext Ultra RNA Library Prep Kit for Illumina following manufacturer’s instructions (NEB, Ipswich, MA, USA). Briefly, mRNAs were first enriched with Oligo(dT) beads. Enriched mRNAs were fragmented for 15 min at 94 °C. First strand and second strand cDNAs were subsequently synthesized. cDNA fragments were end repaired and adenylated at 3’ends, and universal adapters were ligated to cDNA fragments, followed by index addition and library enrichment by limited-cycle PCR. The sequencing libraries were validated on the Agilent TapeStation (Agilent Technologies, Palo Alto, CA, USA), and quantified by using Qubit 2.0 Fluorometer (Invitrogen, Carlsbad, CA) as well as by quantitative PCR (KAPA Biosystems, Wilmington, MA, USA). The sequencing libraries were clustered on 1 lane of a flow cell. After clustering, the flowcell was loaded on the Illumina HiSeq 4000 instrument and the samples were sequenced using a 2×150bp Paired End (PE) configuration. Image analysis and base calling were conducted by the HiSeq Control Software (HCS). Raw sequence data (.bcl files) generated from Illumina HiSeq were converted into fastq files and de-multiplexed using Illumina’s bcl2fastq 2.17 software. One mismatch was allowed for index sequence identification.

Read quality was evaluated using FastQC (*74*) and data were pre-processed with Cutadapt (*75*) for adapter removal following best practices (*76*). Gene expression against the GRCm38 transcriptome (Ensembl 93 version) (*77*) was quantified with STAR (*78*) and featureCounts (*79*). Differential gene expression analysis was performed using DESeq2 (*80*), while ClusterProfiler (*81*) was utilized for functional enrichment investigations. Genes with log2 |Fold Change| ≥1 and False Discovery Rate (FDR) ≤ 0.05 were considered statistically significant.

### Quantitative reverse transcription real-time PCR (qRT-PCR)

The miScript II RT kit (Qiagen) was used according to manufacturer’s protocol for cDNA synthesis from 1 μg of RNA. 30 ng of cDNA were used per PCR reaction. QuantiTect primers were all purchased from Qiagen: *B2m* (QT01149547); *Col1a1* (QT00162204); *Col3a1* (QT01055516); *Egf* (QT00151018); *En1* (QT00248248); *Fgf2* (QT00128135); *Fgf7* (QT00172004); *Fn1* (QT00135758); *Hgf* (QT00158046); *Tgfb1* (QT00145250); *Vegfa* (QT00160769). qRT-PCR was run with QuantiTect SYBR Green PCR kit (Qiagen) on a Stratagene Mx3005P with MxPro qPCR Software (Agilent Technologies). The cycling conditions used were according to the miScript SYBR Green PCR kit protocol (Qiagen). Quantification was performed using the 2–^ΔΔCt^ method. Gene expression was normalized against the housekeeping gene *B2m*.

### Statistical analysis

MATLAB (version R2018b) and Graphpad Prism 7.04 were used to assess the statistical significance of all comparison studies in this work. Data distribution was assumed to be normal for all parametric tests, but not formally tested. In the statistical analysis for comparison between multiple samples, one-way ANOVA followed by Tukey’s or Fisher’s multiple comparison tests were conducted with the threshold of \**p* ≤ 0.05, \*\**p* ≤ 0.01, and \*\*\**p* ≤ 0.001. In the statistical analysis between two data groups, a two-sample Student’s *t*-test was used, and the significance threshold was placed at \**p* ≤ 0.05, \*\**p* ≤ 0.01, and \*\*\**p* ≤0.001.

## Acknowledgments

The authors thank the Koch Institute Swanson Biotechnology Center for technical support, specifically K. Cormier and the Histology Core for the histological processing and analysis.

## Funding

This work is supported by Defense Advanced Research Projects Agency (DARPA) (5(GG0015670)). A.V. received funding from the National Rongxiang Xu Foundation. G.T. received a George and Marie Vergottis Foundation Postdoctoral Fellowship. H.Y. acknowledges the financial support from Samsung Scholarship. H.R. acknowledges the financial support from Kwanjeong Educational Foundation Scholarship.

## Author contributions

G.T., H.Y., A.V., and X.Z. designed the study. H.Y. conceived the idea for the strain-programmable patch. H.Y. and H.R. developed the materials and method for the strain-programmable patch. H.Y., H.R., C.S.N. designed the *in vitro* and *ex vivo* experiment. H.Y. and H.R. conducted the *in vitro* and *ex vivo* experiment and analysis. L.W. and C.F.G. designed and conducted the theoretical and numerical modeling and analysis. J.W. and H.Y. designed and conducted the *in vivo* biocompatibility experiment. G.T., L.C. and A.V. designed and conducted the *in vivo* diabetic wound healing experiment and analysis. G.T., I.M. and N.J. completed the flow cytometry, histology, immunofluorescence, gene expression and human skin *ex vivo* experiment analyses. A.K. performed histology assessment and scoring. X.L.K. and I.S.V. performed sequencing analysis. G.T., H.Y., X.Z., and A.V. wrote the manuscript with inputs from all authors.

## Competing interests

H.Y., H.R., X.Z., G.T., and A.V. are the inventors of a patent application (U.S. Application No. 63/148,901) that covers the design and mechanism of the strain-programmable patch for diabetic wound healing.

## Data and materials availability

All data is available in the main text or the Supplementary Information. The RNA-seq data generated in the present study were deposited in the Gene Expression Omnibus (GEO) with accession number ‘GSE154132‘.

## SUPPLEMENTARY MATERIALS

### Supplementary Texts

#### Hydration-based shape-memory of the strain-programmable patch

To achieve programmable contraction and stress remodeling of wet wounded tissues, we develop a hydration-based shape-memory mechanism and an associated theoretical framework for predictable and reproducible fabrication for the strain-programmable patch. In the following paragraphs, we will discuss (i) a physical picture of the proposed hydration-based shape-memory mechanism, (ii) fabrication of the strain-programmable patch, and (iii) mechanical properties of the strain-programmable patch.

#### 1. Hydration-based shape-memory mechanism

Shape-memory polymers are the class of polymers capable of memorizing a certain macroscopic deformed configuration in the stable fixed state and relaxing to the original undeformed configuration under external stimuli (*28, 29, 82*). The transition from the fixed deformed configuration to the relaxed original configuration relies on a drastic mechanical property changes between the glassy fixed state and the rubbery relaxed state (*28, 29, 82*). In the glassy state, polymer chains in the shape-memory polymers are “frozen” with suppressed mobility and high macroscopic stiffness (i.e., high Young’s modulus). In the rubbery state, polymer chains in the shape-memory polymers recover their entropic elasticity with low macroscopic stiffness (i.e., low Young’s modulus) where the memorized deformation in the fixed glassy state can be elastically relaxed. The glassy to rubbery state transition in shape-memory polymers are conventionally achieved by tuning environmental temperature across the transition temperature (*T*_trans_) where the shape-memory polymers are in the glassy state in low temperature (*T* < *T*_trans_) and in the rubbery state in high temperature (*T* > *T*_trans_), respectively (*27-29, 82*). *T*_trans_ is a critical temperature where the chain mobility of the shape-memory polymers changes drastically, which can be either the glass transition temperature (*T*_g_) for amorphous polymers or the melting temperature (*T*_m_) for crystalline polymers.

In this work, we propose a non-thermal hydration-based shape-memory mechanism without the need of changing environmental temperature to achieve highly programmable and facile contraction and mechanical modulations of wet wounded tissues in synergistic combination with the dry-crosslinking mechanism for rapid robust wet adhesion (*23, 26*). The hydration-based shape-memory mechanism relies on the hydrogels’ unique transition between the glassy state and the rubbery state based on hydration level. In hydrated or swollen state, hydrogels are elastically deformable solids in the rubbery state with low macroscopic stiffness (i.e., low Young’s modulus). However, when hydrogels are dried to remove water, hydrogels transit into the glassy state whose polymer chains are frozen with suppressed mobility and high macroscopic stiffness (i.e., high Young’s modulus), providing ability to stably fix or memorize the deformed configuration and program the applied pre-stretches. By re-hydrating the dried hydrogels, hydrogels can recover their soft and elastic properties in the rubbery state during which the programmed pre-stretches can be elastically relaxed (Fig. 1A).

This hydration-based transition between the glassy and rubber states is due to the reduction of polymer network’s *T*_g_ during hydration. A simplified expression for *T*_g_ is provided as (*24, 25*)

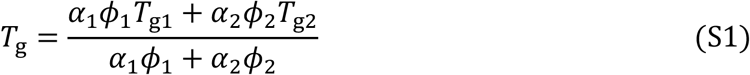

where *α*_1_ and *α*_2_ are the thermal expansion coefficients of the polymer and the solvent, and *ϕ*_1_ and *ϕ*_2_ are the volume fraction of the polymer and solvent, *T*_g1_ and *T*_g2_ are the glass transition temperature of the polymer and the melting point of the solvent, respectively. The melting point *T*_g2_ of water is much lower than the glass transition temperature *T*_g1_ of typical polymers for hydrogels. Notably, the time required to fully hydrate a hydrogel with a thickness of *H* can be expressed as *t*_hydrate_ = (*H*/*k*_h_)^2^ where *k*_h_ is the hydration coefficient of the hydrogel (*26*).

Hence, the proposed hydration-based shape-memory mechanism provides a facile and effective strain-programming and -release strategy for hydrogels, including the bioadhesive layer of the strain-programmable patch, only by natively present water in wet physiological environment without the need of thermal or other complex external stimuli. The hydration-based shape-memory mechanism’s highly versatile and tunable strain-programming capability and water-based biocompatible triggering of the programmed strain release will be particularly advantageous for biomedical and clinical applications operating in wet physiological environments.

#### 2. Fabrication of the strain-programmable patch

Taking advantage of the hydration-based shape-memory mechanism, the strain-programmable patch can be fabricated based on multiple steps of pre-stretch and drying processes as summarized in fig. S3. Since the non-adhesive elastomer backing and the bioadhesive layer in the strain-programmable patch can have different swelling ratio in wet physiological environments (fig. S4), the fabrication process of the strain-programmable patch involves canceling of swelling mismatch between the two layers as well as swelling of the assembled patch.

The fabrication of the strain-programmable patch starts from the as-prepared bioadhesive in the rubbery state with the dimension of 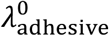 (=1.48 for the bioadhesive used in this work) in length and width and 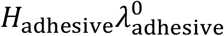 in thickness where the reference state (i.e., isotropically dried bioadhesive) has the dimension of *H*_adhesive_ in thickness and unity in length and width. The fabrication process consists of five distinctive steps as the following:

##### Step 1

Introduce the non-adhesive elastomer backing resin (i.e., before curing) on the as-prepared bioadhesive layer, and pre-stretch the bioadhesive layer by ratio of

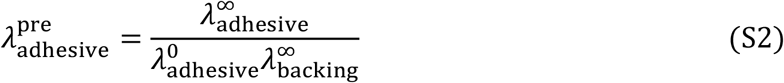

in both directions to cancel out the swelling mismatch between the non-adhesive elastomer backing and the bioadhesive layer (Step 1 in fig. S3), where 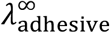 is the equilibrium swelling ratio of the bioadhesive layer and 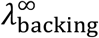 is the equilibrium swelling ratio of the non-adhesive elastomer backing. For the bioadhesive and the elastomer backing used in this work, 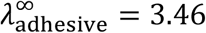 and 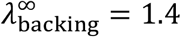 which give 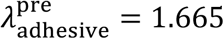 (fig. S4).

After this step, the bioadhesive layer in the rubbery state has the dimension of 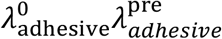 in length and width and 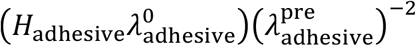 in thickness.

##### Step 2

Cure the non-adhesive elastomer backing on the pre-stretched bioadhesive layer (Step 2 in fig. S3).

After this step, the non-adhesive elastomer backing in the rubbery state has the dimension of 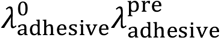 in length and width and *H*_backing_ in thickness; the bioadhesive layer in the rubbery state has the dimension of 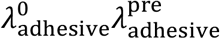 in length and width and 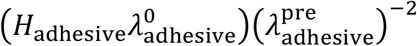 in thickness.

##### Step 3

Pre-stretch both non-adhesive elastomer backing and bioadhesive layer by ratio of

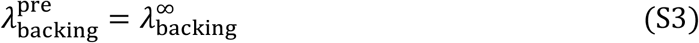

in both directions to cancel out the dimensional change of the patch by swelling in wet physiological environments (Step 3 in fig. S3).

After this step, the non-adhesive elastomer backing in the rubbery state has the dimension of 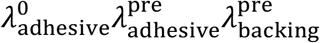 in length and width and 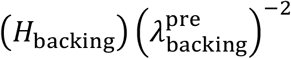 in thickness; the bioadhesive layer in the rubbery state has the dimension of 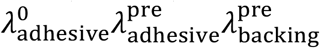 in length and width and 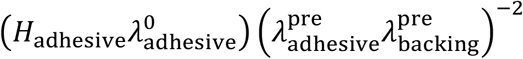 in thickness.

##### Step 4

Pre-stretch both non-adhesive elastomer backing and bioadhesive layer by ratios of 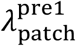 and 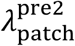 to each direction to program desired strains (direction 1 in length, direction 2 in width), respectively (Step 4 in fig. S3).

After this step, the non-adhesive elastomer backing in the rubbery state has the dimension of 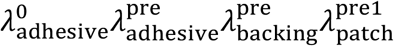 in length, 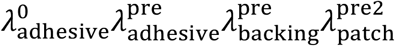 in width, and 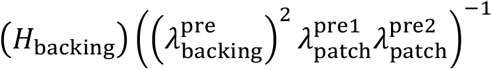 in thickness; the bioadhesive layer in the rubbery state has the dimension of 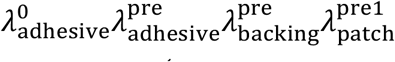 in length, 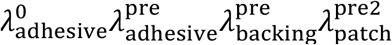 in width, and 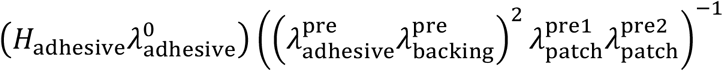 in thickness.

##### Step 5

Dry the assembled strain-programmable patch to shape-memory the pre-stretched configuration (Step 5 in fig. S3), completing the fabrication of the strain-programmable patch.

After this step, the non-adhesive elastomer backing in the rubbery state has the dimension of 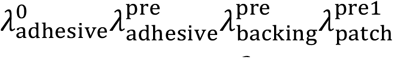 in length, 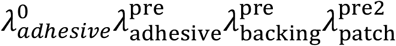 in width, and 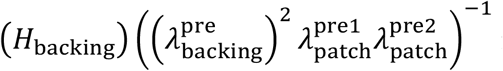 in thickness; the bioadhesive layer in the glassy state has the dimension of 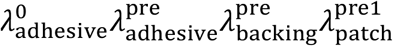 in length, 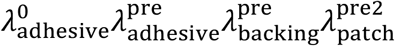 in width, and 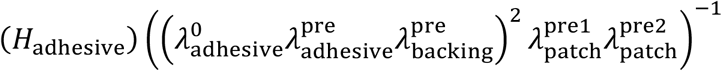 in thickness.

#### 3. Mechanical properties of the strain-programmable patch

##### 3.1. Measurement of physical parameters of the strain-programmable patch

Since both non-adhesive elastomer backing and bioadhesive layer of the strain-programmable patch used in this work become hydrogel in wet physiological environments, we take the swollen strain-programmable patch as a Flory-Rehner hydrogel with thermodynamic parameters of *N*_adhesive_, *N*_backing_, *χ*_adhesive_, and *χ*_backing_ for simplicity of the analysis, where *N*_adhesive_ is the number of polymer chains per unit volume of the bioadhesive at the reference state (fig. S3), *N*_backing_ is the number of polymer chains per unit volume of the elastomer backing at the reference state (fig. S3), *χ*_adhesive_ is the Flory solvent-polymer interaction parameter for the bioadhesive, and *χ*_backing_ is the Flory solvent-polymer interaction parameter for the elastomer backing (*83, 84*).

To determine *N*_adhesive_, the shear modulus 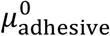 is measured for the as-prepared bioadhesive and *N*_adhesive_ is then calculated from (*85*)

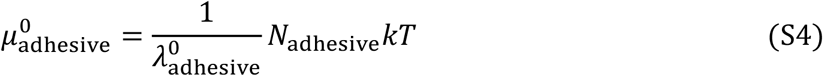

where *k* is the Boltzmann constant, *T* is the absolute temperature. To measure *χ*_adhesive_, we carry out free-swelling experiment for the as-prepared bioadhesive without constraint (i.e., freely swelling in PBS). The Cauchy stress *σ*_eq,adhesive_ in any direction of the unconstrained equilibrium swollen bioadhesive can be expressed as (*85*)

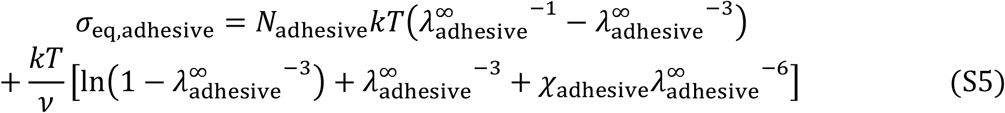

where *v* is the volume of the water molecule. Due to the traction-free boundary condition, the Cauchy stress *σ*_eq,adhesive_ is zero in Eq. S5. In this study, we used *T* = 310 K (i.e., body temperature), *v* = 3.0 × 10^−29^ m^3^ and experimentally-measured values of 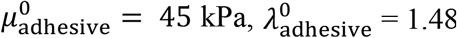, and 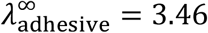. By implementing these values and solving Eq. S5, we can calculate *N*_adhesive_ = 1.56 × 10^25^ m^−3^ and *χ*_adhesive_ = 0.29.

Similarly, to determine *N*_backing_, the shear modulus 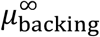 is measured for the equilibrium swollen elastomer backing and *N*_backing_ is then calculated from (*85*)

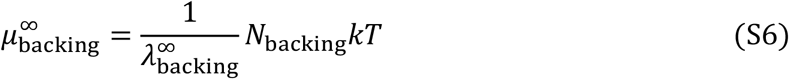

To measure *χ*_backing_, we carry out free-swelling experiment for the elastomer backing without constraint (i.e., freely swelling in PBS). The Cauchy stress *σ*_eq,backing_ in any direction of the unconstrained equilibrium swollen elastomer backing can be expressed as (*85*)

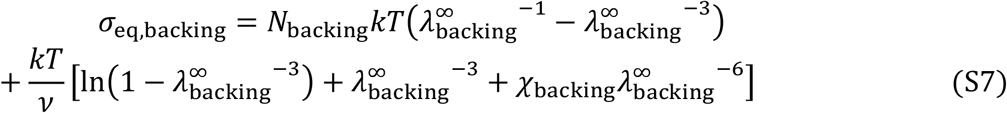

Due to the traction-free boundary condition, the Cauchy stress *σ*_eq,backing_ is zero in Eq. S7. In this study, we used *T* = 310 K (i.e., body temperature), *v* = 3.0 × 10^−29^ m^3^ and experimentally-measured values of 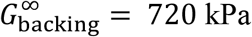 and 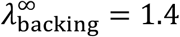. By implementing these values and solving Eq. S7, we can calculate *N*_backing_ = 2.36 × 10^26^ m^3^ and *χ*_backing_ = 0.65.

##### 3.2. Flory-Rehner model of the strain-programmable patch

In the equilibrium swollen state, the Cauchy stresses (or true stresses) generated by the bioadhesive layer *σ*_adhesive_ and the elastomer backing *σ*_backing_ can be expressed as (*85, 86*)

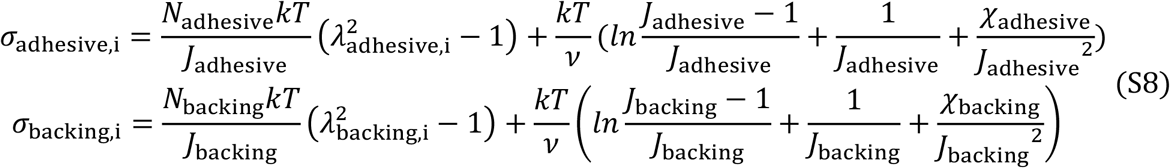

where *σ*_i_ is the Cauchy stress for each principal direction (i = 1 for length, 2 for width, and 3 for thickness direction, respectively), *λ*_i_ is the stretch from the reference state for each principal direction (i = 1 for length, 2 for width, and 3 for thickness direction, respectively), *J*_adhesive_ = *λ*_adhesive,1_*λ*_adhesive,2_*λ*_adhesive,3_, and *J*_backing_ = *λ*_backing,1_*λ*_backing,2_*λ*_backing,3_.

Since the strain-programmable patch consists of the elastomer backing and the bioadhesive layer, the overall Cauchy stress generated by the strain-programable patch in equilibrium swollen state can be expressed as

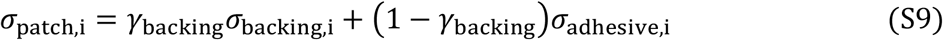

where *γ*_backing_ is the ratio of the elastomer backing thickness in the total thickness of the equilibrium swollen strain-programmable patch which can be calculated as

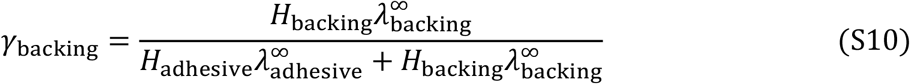

where *H*_backing_ and *H*_adhesive_ are thicknesses of the elastomer backing and the bioadhesive at the reference state, respectively (fig. S3).

##### 3.3. Neo-Hookean model approximation of the strain-programmable patch

While we model the strain-programmable patch in equilibrium swollen state as a Flory-Rehner hydrogel, we also investigate approximation of the strain-programmable patch as an incompressible neo-Hookean solid (*87*) for simplicity of the analysis, particularly in finite-element modeling. To obtain shear moduli of materials for incompressible neo-Hookean models, we first fit the uniaxial tensile test data of the elastomer backing, the bioadhesive layer, and the strain-programmable patch in equilibrium swollen state by using the following expression

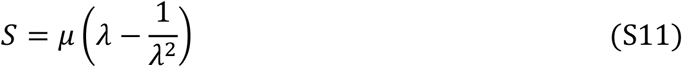

where *S* is the nominal or engineering stress, *µ* is the shear modulus, and *λ* is the stretch of sample in tensile tests. As a result, we obtain shear moduli of the elastomer backing 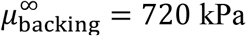, the bioadhesive layer 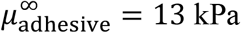, and the strain-programmable patch 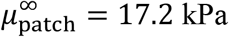 in equilibrium swollen state, respecitvely (Fig. 2D and fig. S2).

As illustrated in fig. S5, the Flory-Rehner model and the incompressible neo-Hookean model show good agreement for the equilibrium swollen strain-programmable patch. This indicates that the simpler incompressible neo-Hookean model can be used to describe the mechanical behavior of the strain-programmable patch.

### Modeling of wound contraction and stress remodeling by the strain-programmable patch

To provide quantitative and predictive design guideline for contraction and stress remodeling of wounds by the strain-programmable patch, we develop analytical and finite-element models based on mechanical properties of the strain-programmable patch and tissues. In this study, without loss of generality, our models are developed based on a circular wound in the skin which is a common form for diabetic wounds and an equibiaxially strain-programmed patch (i.e., 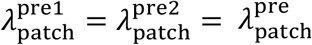).

#### 1. Analytical modeling

##### 1.1. Initial enlarging of wound by pre-strain in native skin

A skin wound with the initial radius denoted as *a* enlarges to *a*_1_immediately after wounding due to the relaxation of the pretension *σ*_∞_ around the wound edge generated by the pre-strain in native skin. The reference state refers to an imaginary stress-free state in which 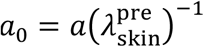 where 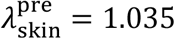 is the experimentally measured pre-strain within the native skin. When the strain-programmable patch is applied to the wounded skin, the wound shrinks to radius *a*_closure_ (fig. S12). Axisymmetric condition is adopted for both skin and patch for equibiaxial scenario and the outer boundary of skin is assumed to be much larger than the wound (i.e., 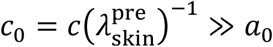).

The strain-programmable patch initially has a radius of *b*_0_ and undergoes an equibiaxial stretch of 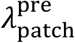. For an incompressible neo-Hookean material under equibiaxial stretch, the Cauchy stresses are calculated as

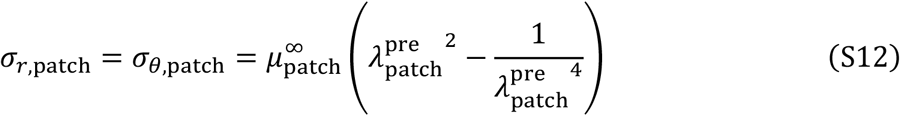

where 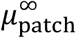 is the shear modulus of the strain-programmable patch in equilibrium swollen state. The skin is modeled as an incompressible Ogden material and the pretension in the skin before wounding can be found as

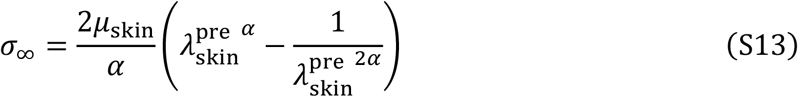

where *μ*_skin_ = 40 kPa and α = 20 (for human skin) are fitting parameters measured from experiment. By plugging 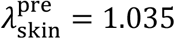 into Eq. S13, one can calculate that *σ*_∞_ = 0.17*μ*_skin_.

Next, we can solve the wound radius *a*_1_ immediately after wounding of the skin. Consider an arbitrary circle with radius *R* ≥ *a*_0_ in the reference state and moves to *r* in the deformed configuration. The incompressibility requires

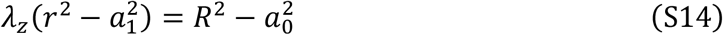

where *λ*_*z*_ is the stretch in z-direction. Then

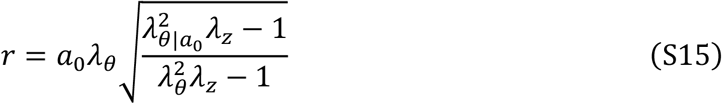

where hoop stretch *λ*_*θ*_ = *r*/*R* and 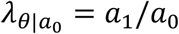 is the hoop stretch at the wound boundary *R* = *a*_0_ and *λ*_*r*_*λ*_*θ*_*λ*_*z*_ = 1. The equilibrium equation in current configuration can be expressed as

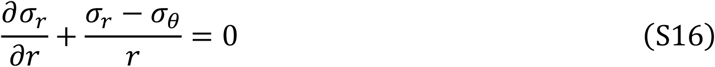

To solve this, we can change variable from *r* to *λ*_*θ*_. With Eq. S16, we have

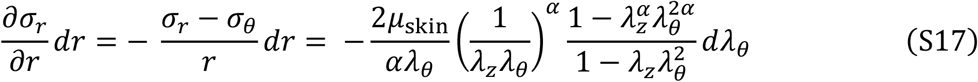

By integrating Eq. S17 from *r* = *a* to arbitrary *r* > *a*_1_, we obtain

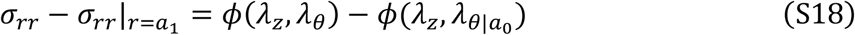

where *ϕ*(*λ*_*z*_, *λ*_*θ*_) is analytical function of *λ*_*z*_, *λ*_*θ*_ and 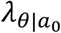 is the hoop stretch at *r* = *a*_0_. Due to the lengthy expression, we do not provide the complete form of *ϕ*(*λ*_*z*_, *λ*_*θ*_) here. By invoking the boundary conditions at 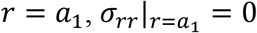, and at 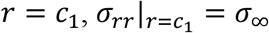, one can have

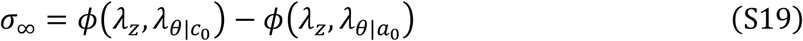

where 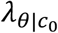 is the hoop stretch at *r* = *c*_0_. Also, from Eq. S18, the hydrostatic pressure in 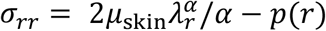 can be solved as

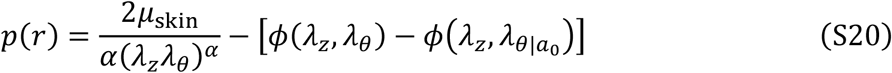

Therefore, the Cauchy stress in the z-direction can be expressed as

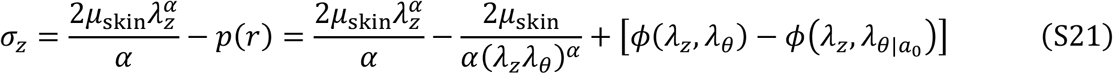

To solve Eq. S21, instead of assuming *σ*_*z*_ = 0, we adopt the relaxed boundary condition of zero resultant force at the arbitrary z-plane (*88*) as

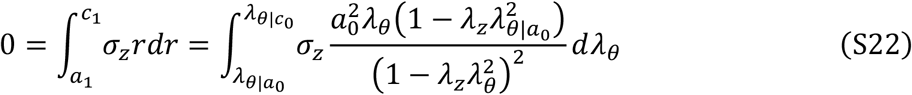

where the incompressibility enforces that

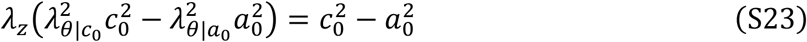

Therefore, one can solve three unknowns 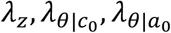 with Eqs. S19, S22, and S23. Then, we can solve the deformation field

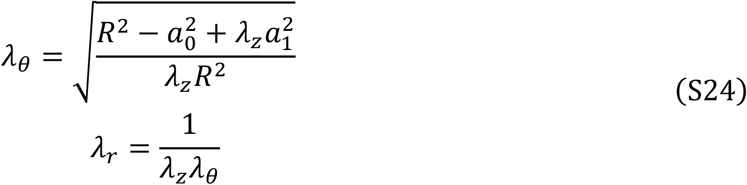

as well as the stress field

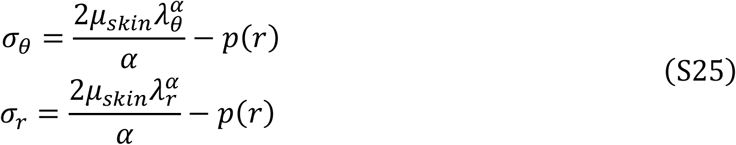

which are plotted in fig. S13. It is apparent that when the out boundary is considerably large (i.e., *c*_0_/*a*_0_ ≫ 1), the converged analytical solutions show that hoop stretch at wound edge 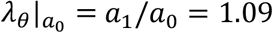. In other words, the size of wound becomes 1.09 times larger than the initial size (i.e., *a*_1_/*a* = 1.05) due to the presence of pre-strain in the native skin.

##### 1.2. Contraction of wound by the strain-programmable patch

Next, we can solve *a*_closure_ after applying the strain-programmable patch to the wounded skin. Analytical solutions can be obtained when the strain-programmable patch has the same size as that of the enlarged wound (i.e., 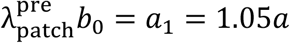) without overlapping area. Let 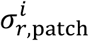 and 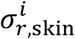 denote the interfacial radial stress on the strain-programmable patch and the skin, respectively. The force balance requires that

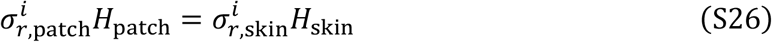

where *H*_patch_ and *H*_skin_ are thickness of the strain-programmable patch and the skin, respectively. As for the skin, stress boundary conditions at *r* = *a*_closure_ now changes to 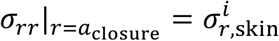. Similar analysis can be performed to find three equations as

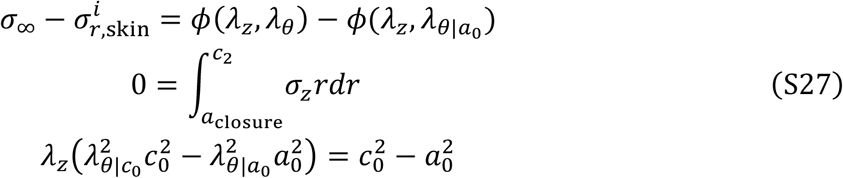

from which one can solve 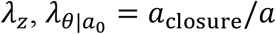, and 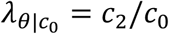 as a function of 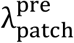. The analytical solution from Eq. S27 are plotted in Figures 4D and S15D which validates the finite-element results. Note that the analytical solutions are only available when the strain-programmable patch size equals to *a*_1_ = 1.1*a* for diabetic mouse skin and *a*_1_ = 1.05*a* for human skin. For the strain-programmable patch with larger sizes, finite-element modeling-based analysis is required as discussed in the following section.

#### 2. Finite-element modeling

To quantitatively analyze the closure and stress remolding of wounds by the strain-programmable patch larger than the wound size (i.e., *b*_0_ > 1.1*a* for diabetic mouse skin, *b*_0_ > 1.05*a* for human skin), we develop 2D axisymmetric finite-element models based on a commercially-available software (ABAQUS/Standard 2017, Dassault Systèmes^®^) for both diabetic mouse skin and porcine skin. The finite-element setups are illustrated in fig. S14. The diabetic mouse skin was modeled as an incompressible Ogden hyperelastic solid (fitting parameters: *μ*_mouse_ = 87.5 kPa, *α*_mouse_ = 7) with 3.5 % tensile pre-strain. The human skin was modeled as an incompressible Ogden hyperelastic solid (fitting parameters: *μ*_human_ = 40 kPa, *α*_human_ = 20) with 3.5 % tensile pre-strain (*36*). The strain-programmable patch was modeled as a neo-Hookean solid (fitting parameter: 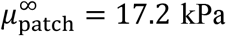) with varying patch sizes and pre-stretches for strain-programming. Three mechanical quantities, (i) wound closure ratio defined as *a*_closure_/*a* (Fig. 4D and fig. S15D), (ii) normalized hoop stress *σ*_*θ*_/*σ*_∞_ (Fig. 4E and fig. S15E), and (iii) normalized radial stress *σ*_*r*_/*σ*_∞_ (Fig. 4F and fig. S15F) are obtained from the finite-element models for both diabetic mouse skin and human skin.

## Supplementary Figures

**Fig. S1.**
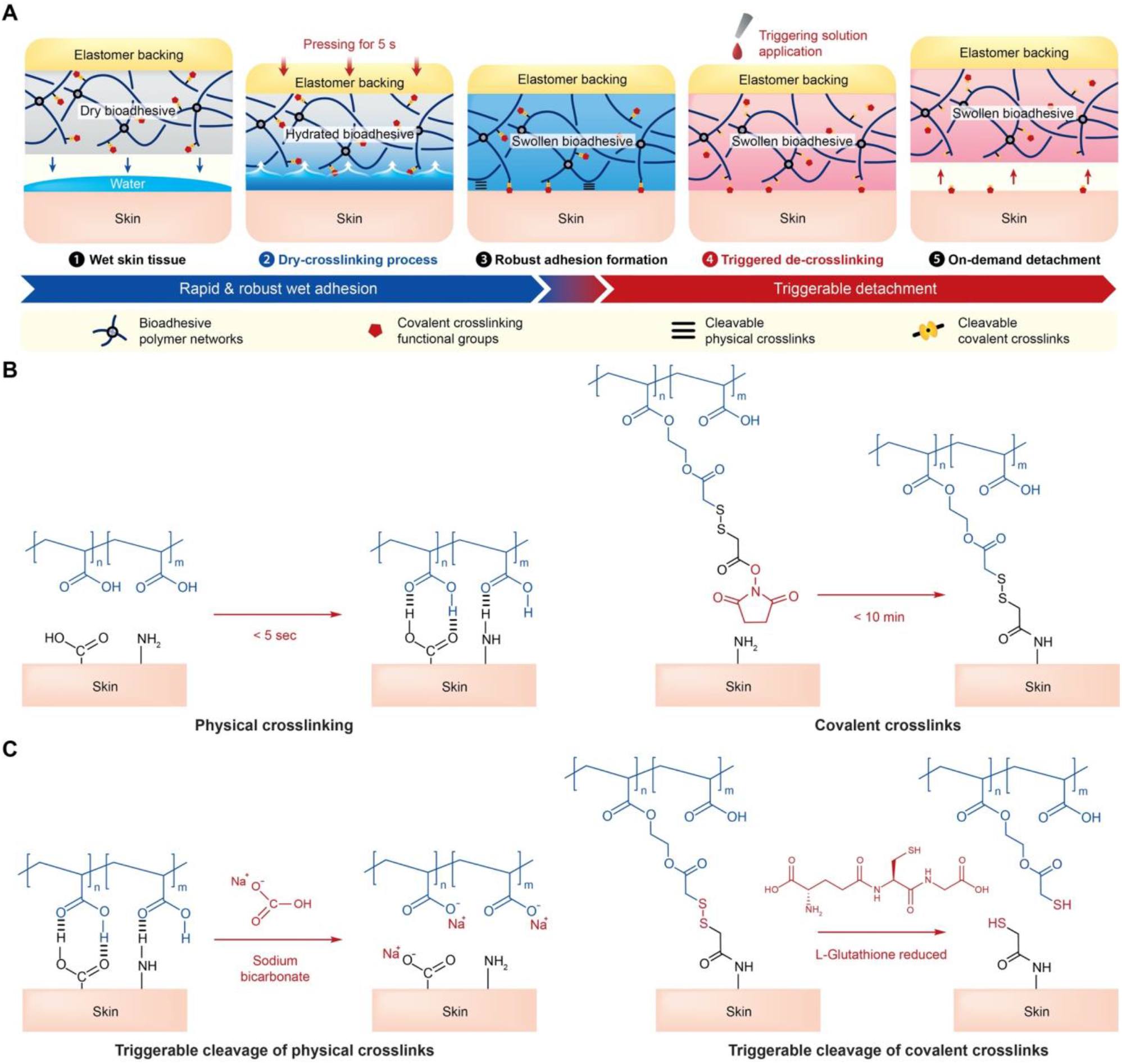
Rapid wet adhesion and triggerable detachment of a strain-programmable patch. (**A**) Schematics illustration for rapid wet adhesion and triggerable detachment of the strain-programmable patch. (**B**) Chemistry of physical and covalent crosslinks for rapid wet adhesion. (**C**) Chemistry of on-demand triggerable detachment.

**Fig. S2.**
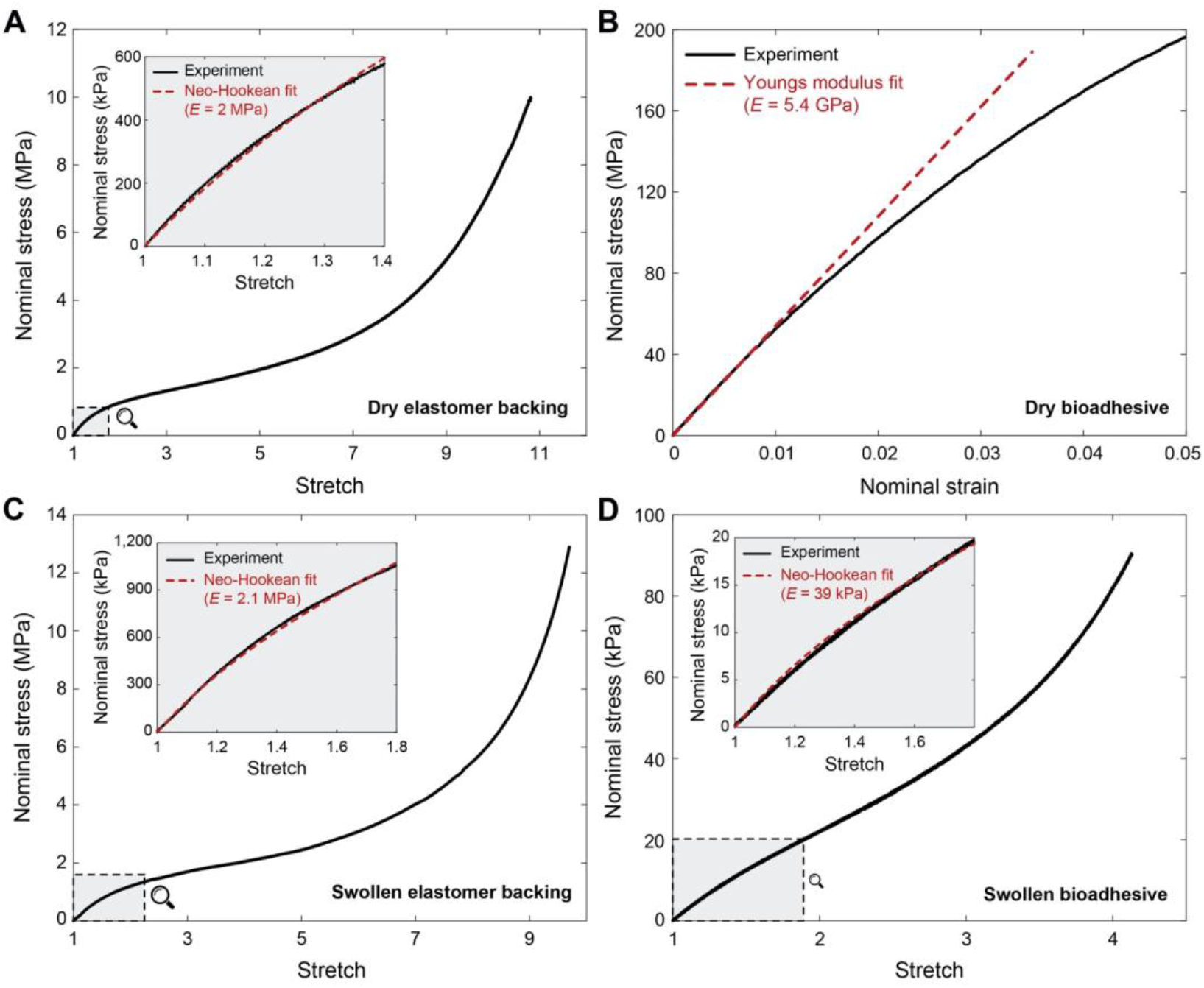
Mechanical properties of elastomer backing and bioadhesive. (**A** and **B**) Nominal stress vs. stretch curves for dry elastomer backing (A) and bioadhesive (B). (**C** and **D**) Nominal stress vs. stretch curves for swollen elastomer backing (C) and bioadhesive (D).

**Fig. S3.**
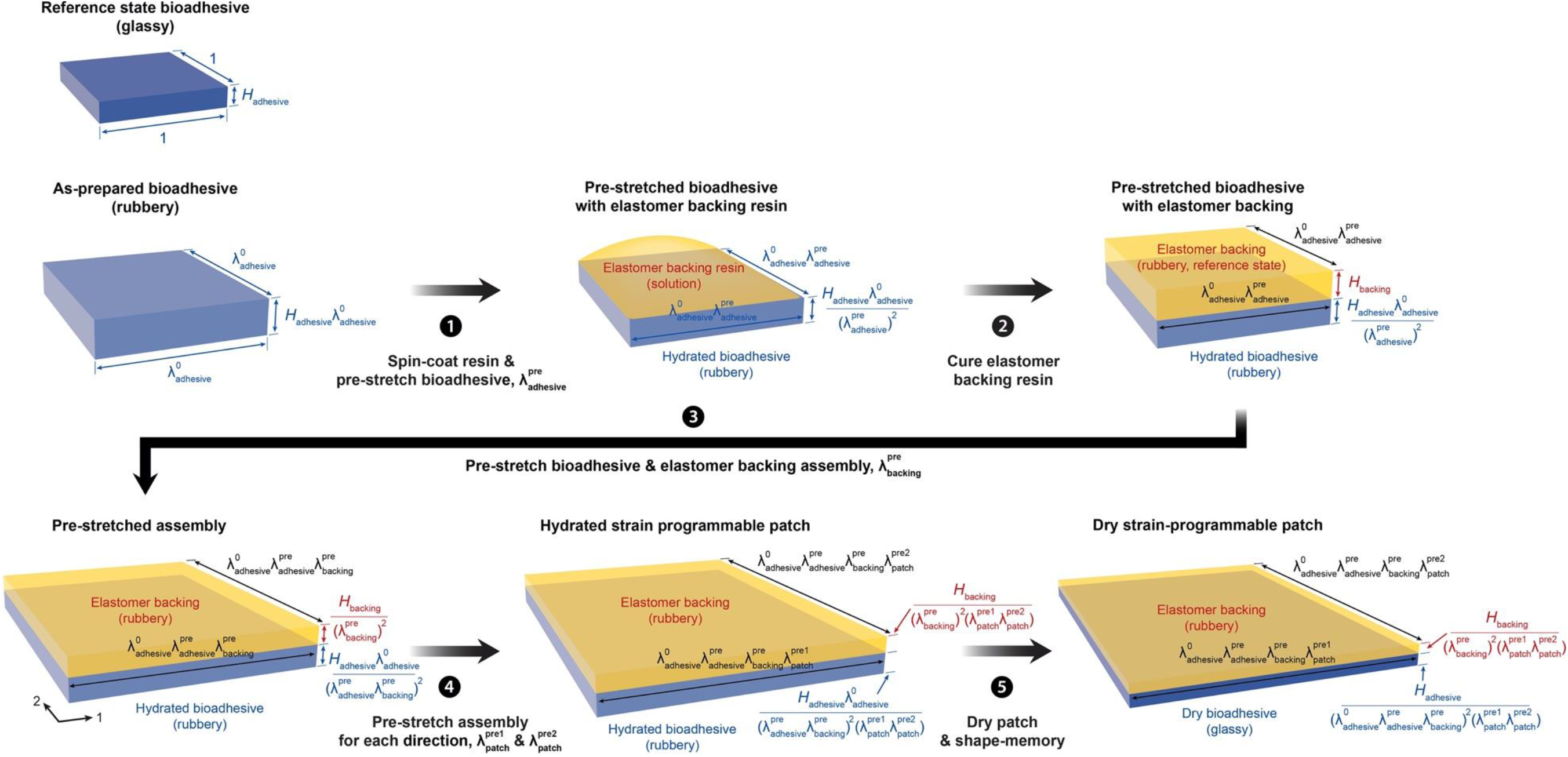
Fabrication of the strain-programmable patch. (1) Spin-coat elastomer backing resin on an as-prepared bioadhesive and pre-stretch the as-prepared bioadhesive by ratio of 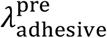. (2) Cure elastomer backing resin while keeping the bioadhesive hydrated. (3) Pre-stretch the bioadhesive and elastomer backing assembly by ratio of 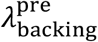. (4) Pre-stretch the bioadhesive patch along two in-plane directions by ratios of 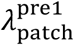 and 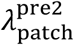, respectively. (5) Dry the bioadhesive for shape-memory to finalize fabrication of the strain-programmed patch.

**Fig. S4.**
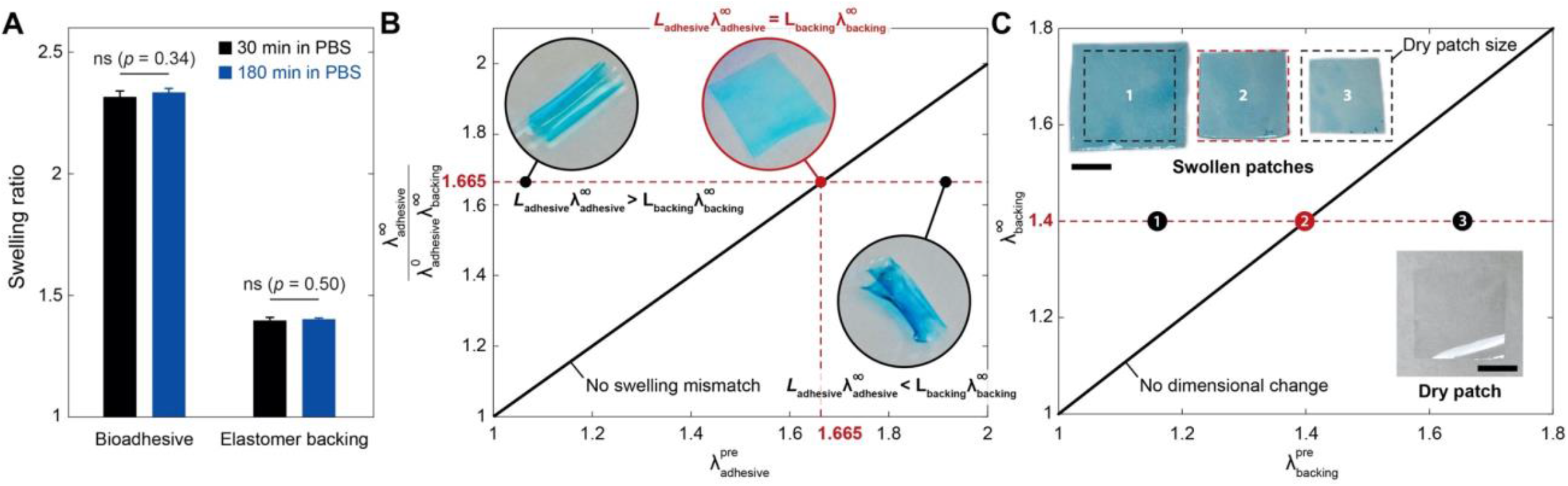
Swelling of strain-programmable patch. (**A**) Swelling ratios of elastomer backing and bioadhesive in wet physiological environment. (**B**) Swelling mismatch canceling between elastomer backing and bioadhesive in the strain-programmable patch. (**C**) Swelling canceling of the strain-programmable patch. Values in A represent the means ± SD (*n* = 4). *P* values are determined by a Student’s *t* test; ns, not significant. Scale bars, 10 mm (C).

**Fig. S5.**
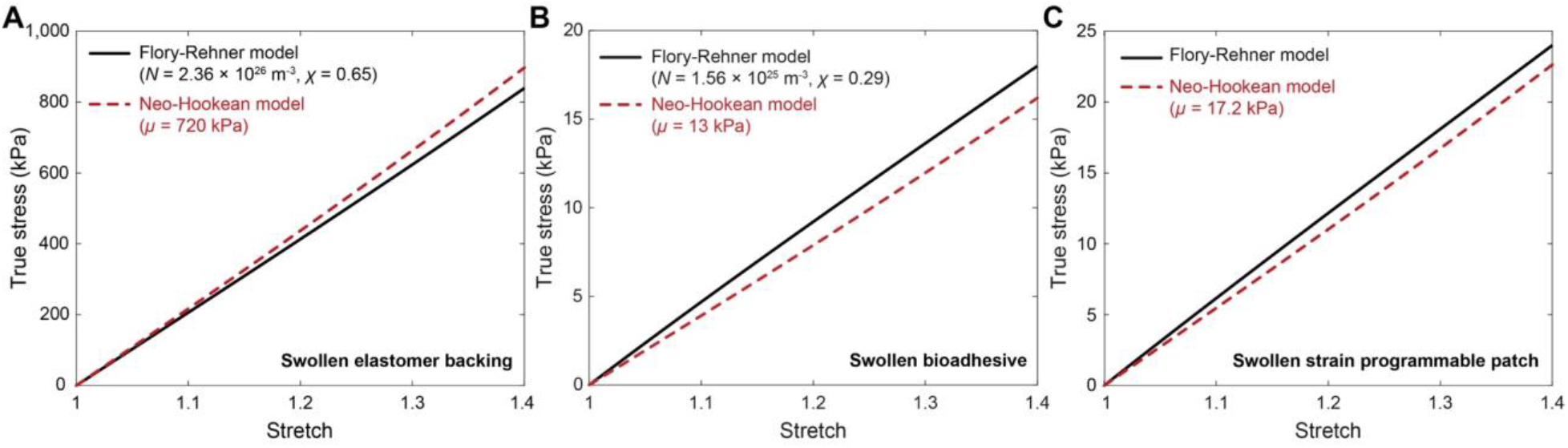
Comparison between Flory-Rehner and neo-Hookean models. (**A** to **C**) True stress vs. stretch obtained based on Flory-Rehner and neo-Hookean models for swollen elastomer backing (A), bioadhesive (B), and strain-programmable patch (C).

**Fig. S6.**
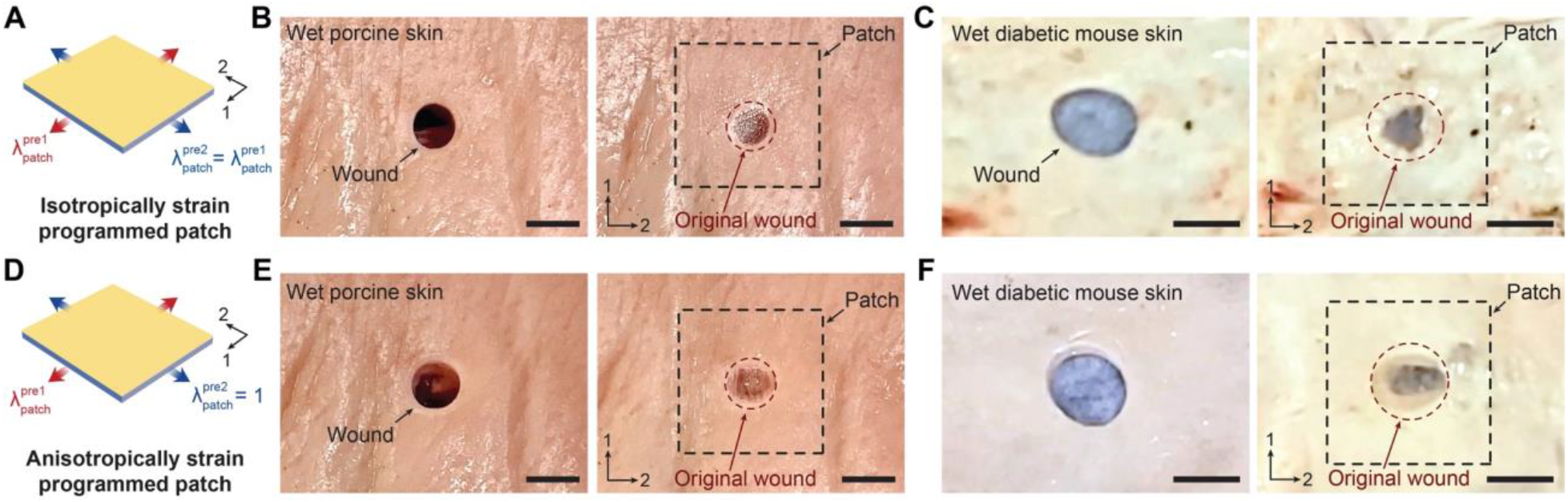
Isotropic and anisotropically strain-programming of the patch. (**A**) Schematic illustration for the isotropically strain-programmed patch. (**B** and **C**) Isotropically strain-programmed patches on a porcine skin (B) and a diabetic mouse skin (C) with circular wound. (**D**) Schematic illustration for the anisotropically strain-programmed patch. (**E** and **F**) Anisotropically strain-programmed patches on a porcine skin (E) and a diabetic mouse skin (F) with circular wound. Scale bars, 10 mm (B and E); and 5 mm (C and F).

**Fig. S7.**
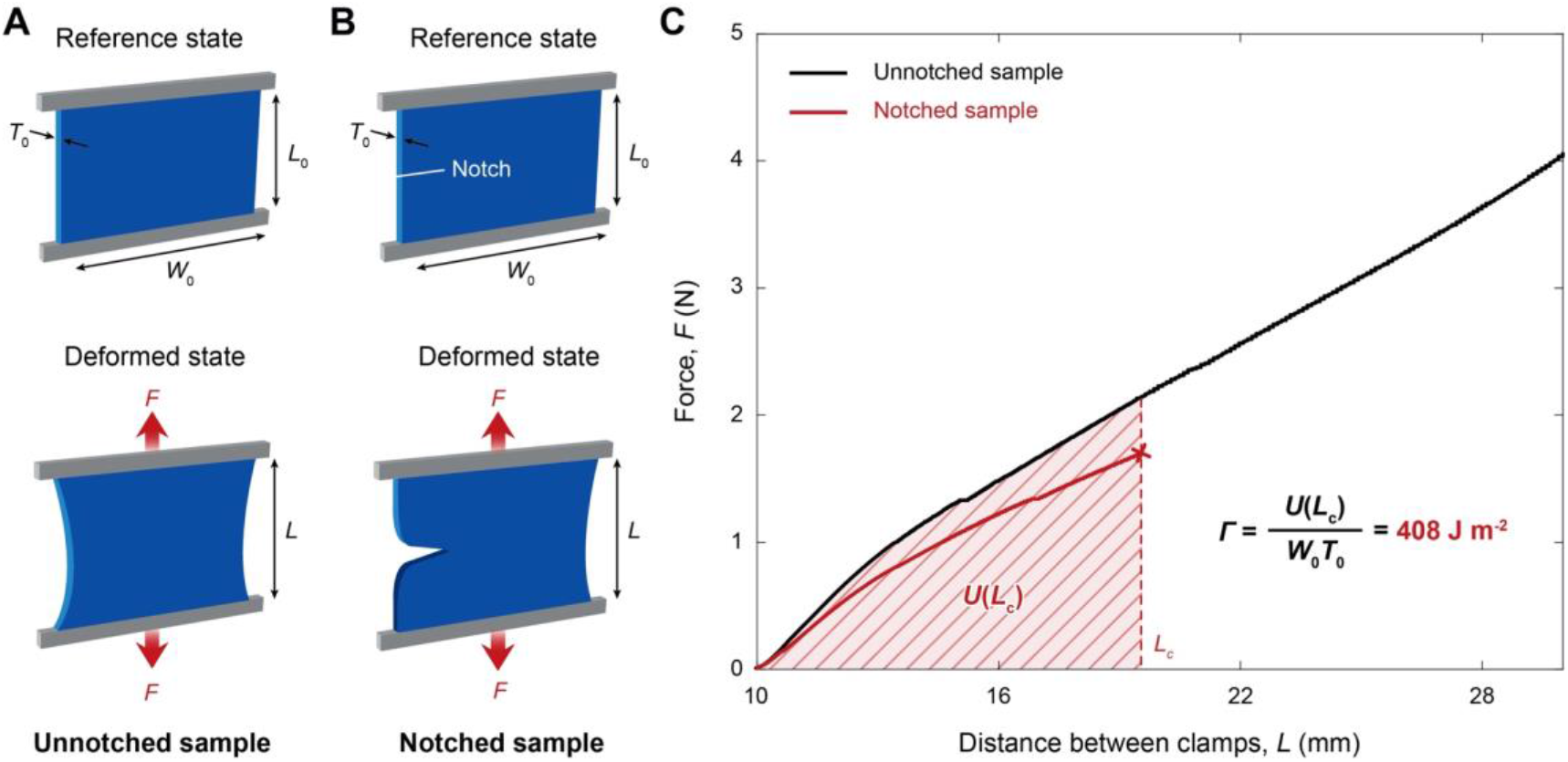
Fracture toughness of strain-programmable patch. (**A** and **B**) Schematic illustrations of pure-shear test for an unnotched sample (A) and a notched sample (B). (**C**) Force vs. distance between clamps for the unnotched and notched swollen strain-programmable patch for fracture toughness measurement. *L*_c_ indicates the critical distance between the clamps at which the notch turns into a running crack. The measured fracture toughness of the strain-programmable patch is 408 J m^-2^.

**Fig. S8.**
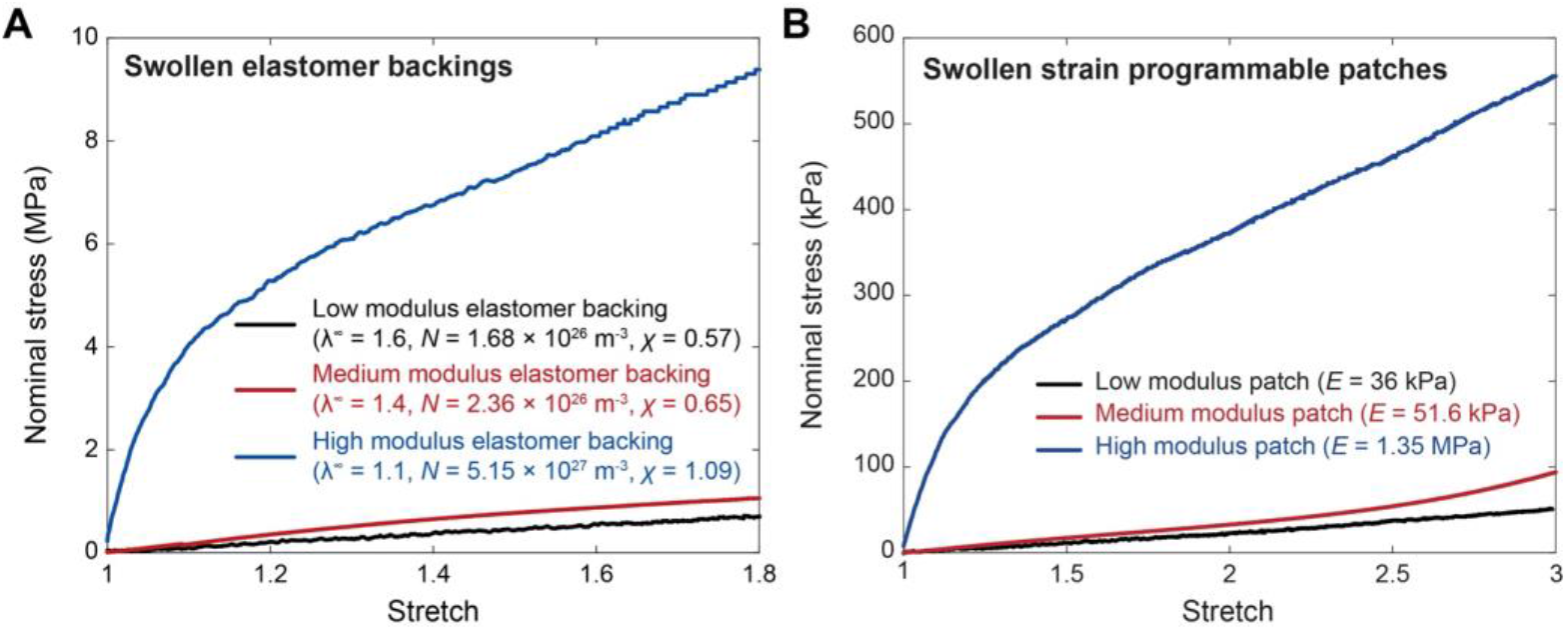
Various elastomer backings for the strain-programmable patch. (**A**) Nominal stress vs. stretch curves for various elastomer backing materials. (**B**) Nominal stress vs. stretch curves for the strain-programmable patch with various elastomer backing materials.

**Fig. S9.**
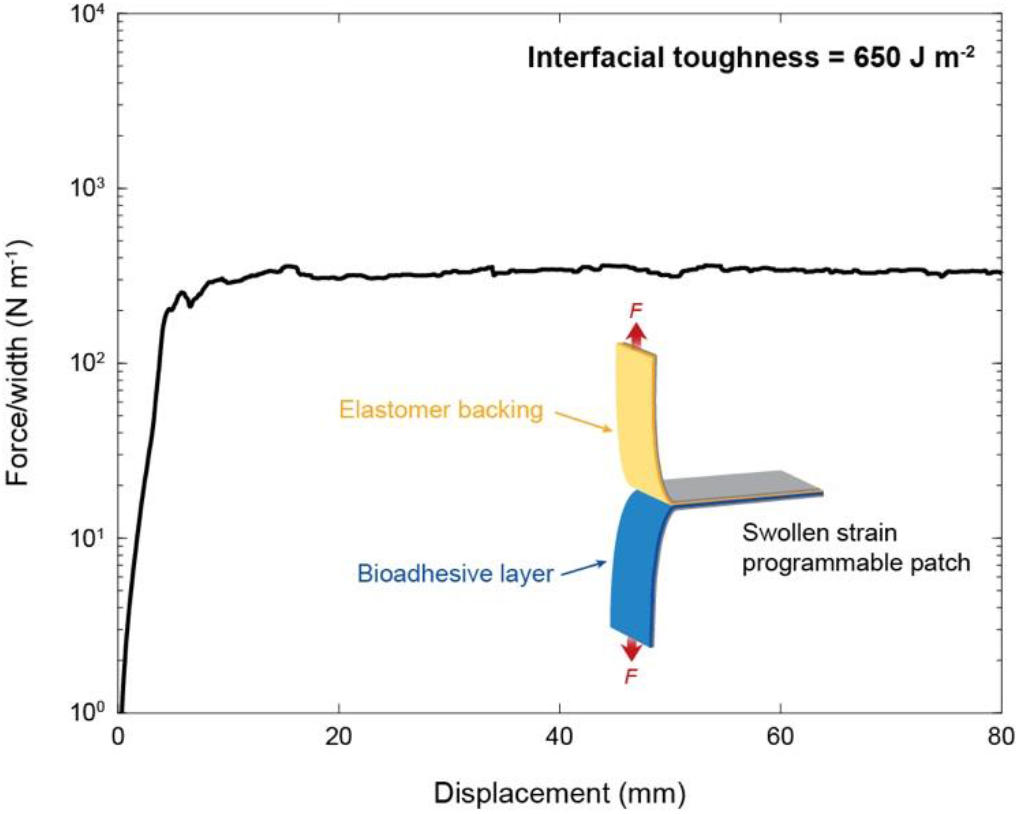
Interfacial toughness between elastomer backing and bioadhesive in the strain-programmable patch. The measured interfacial toughness between swollen elastomer backing and bioadhesive is 650 J m^-2^.

**Fig. S10.**
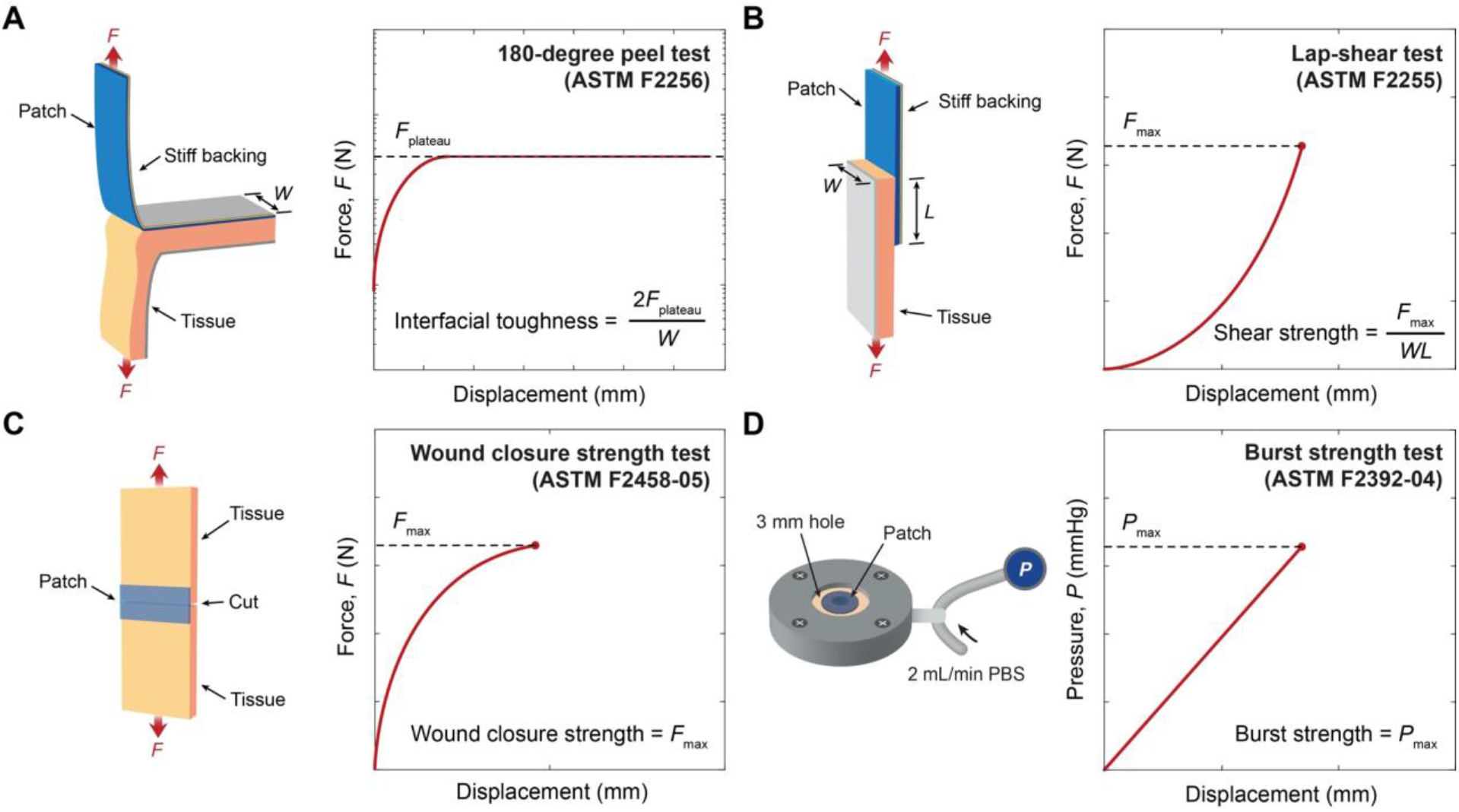
Mechanical testing setups for evaluation of adhesion performance. (**A**) Testing setup for interfacial toughness measurements based on the standard 180-degree peel test (ASTM F2256). **(B)** Testing setup for shear strength measurements based on the standard lap-shear test (ASTM F2255). (**C**) Testing setup for wound closure strength measurements based on the standard tensile test (ASTM F2458-05). (**D**) Testing setup for burst strength measurements based on the standard burst pressure test (ASTM F2392-04).

**Fig. S11.**
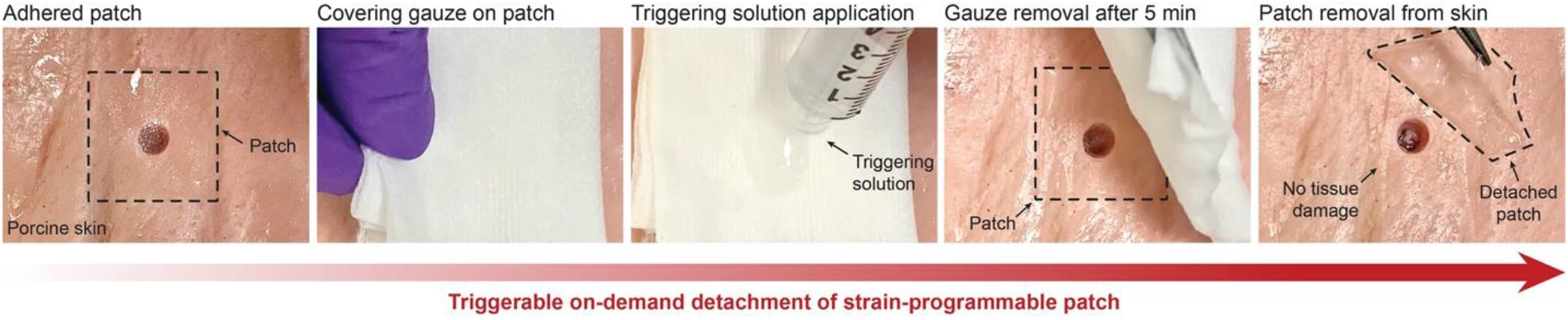
Triggerable on-demand detachment of the strain-programmable patch.

**Fig. S12.**
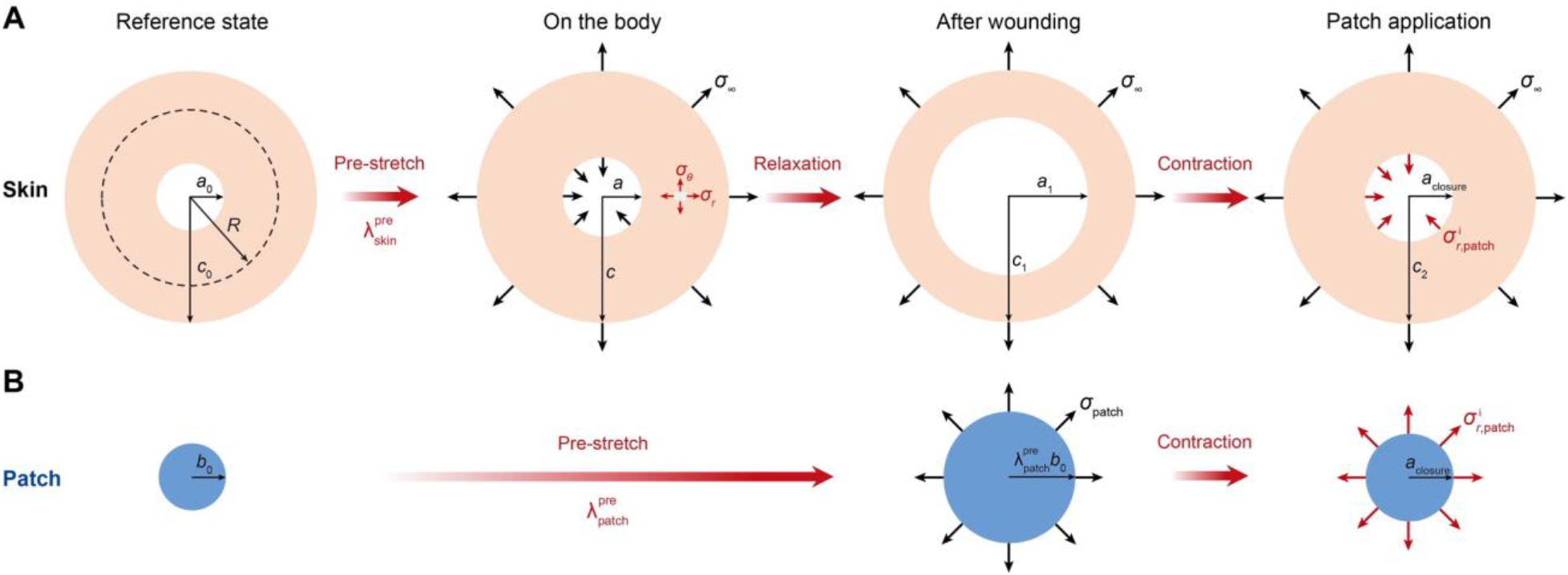
Schematic configuration for the analytical modeling. (**A** and **B**) Axisymmetric configuration of the wounded skin (A) and the strain-programmable patch (B).

**Fig. S13.**
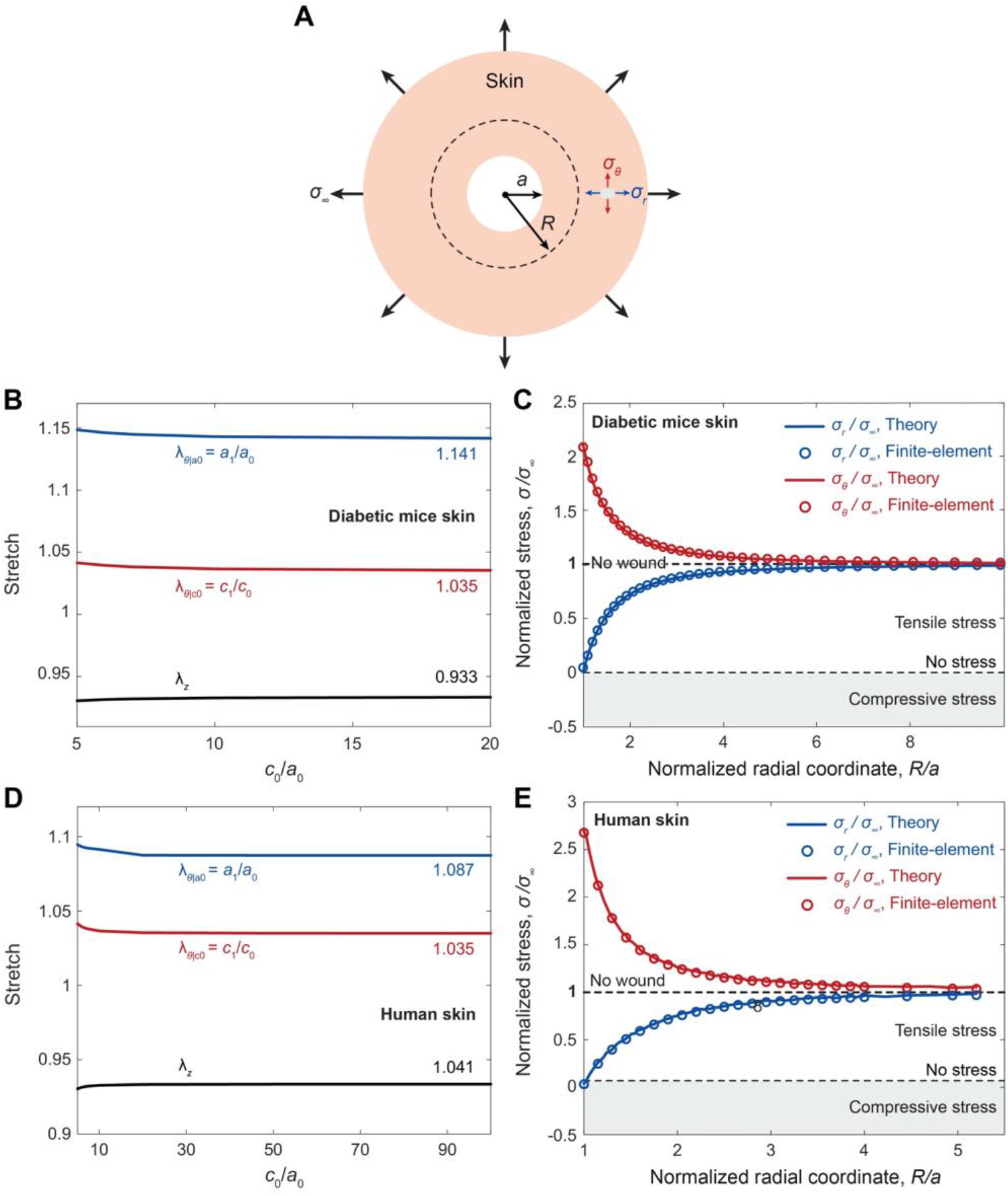
Comparison of analytical and finite-element analyses. (**A**) Schematic illustration for axisymmetric configuration for analytical and finite-element analyses. (**B**) Analytically solved stretches of the wounded diabetic mouse skin as a function of *c*_0_/*a*_0_. (**C**) Stress distribution within the wounded diabetic mouse skin calculated from the analytical solutions (solid lines) and the corresponding finite-element results (circles). (**D**) Analytically solved stretches of the wounded human skin as a function of *c*_0_/*a*_0_. (**E**) Stress distribution within the wounded human skin calculated from the analytical solutions (solid lines) and the corresponding finite-element results (circles).

**Fig. S14.**
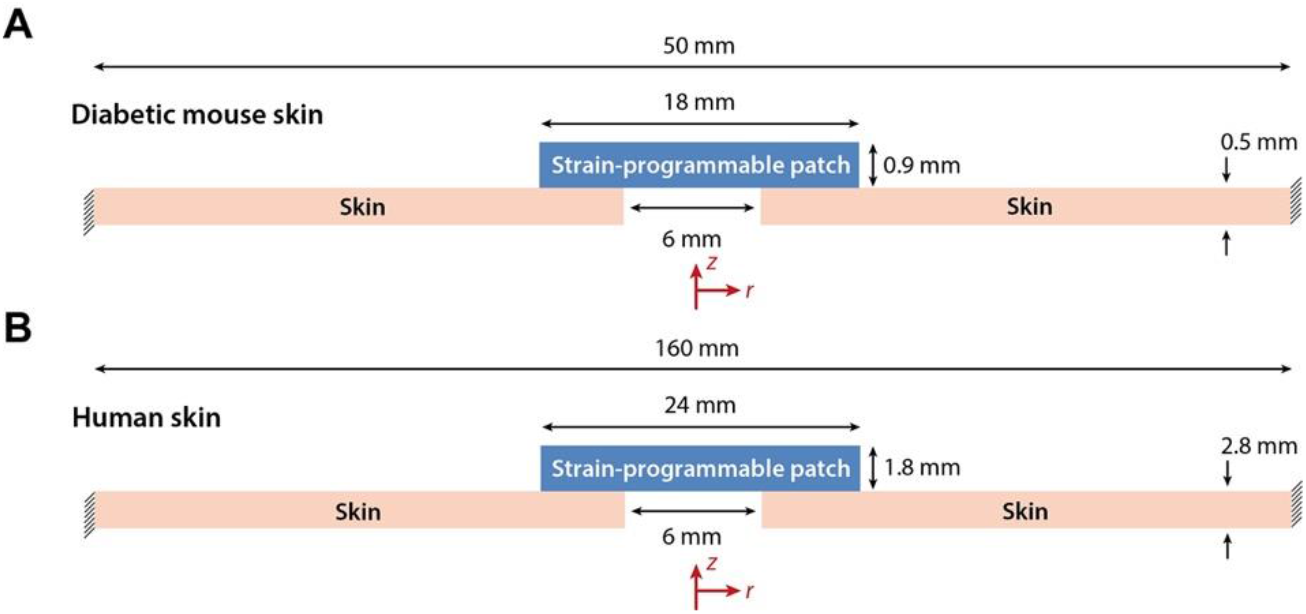
Finite-element modeling for wound closure by the strain-programmable patch. (**A** and **B**) Finite-element setups for diabetic mouse skin (A) and human skin (B) with wound and the strain-programmed patch.

**Fig. S15.**
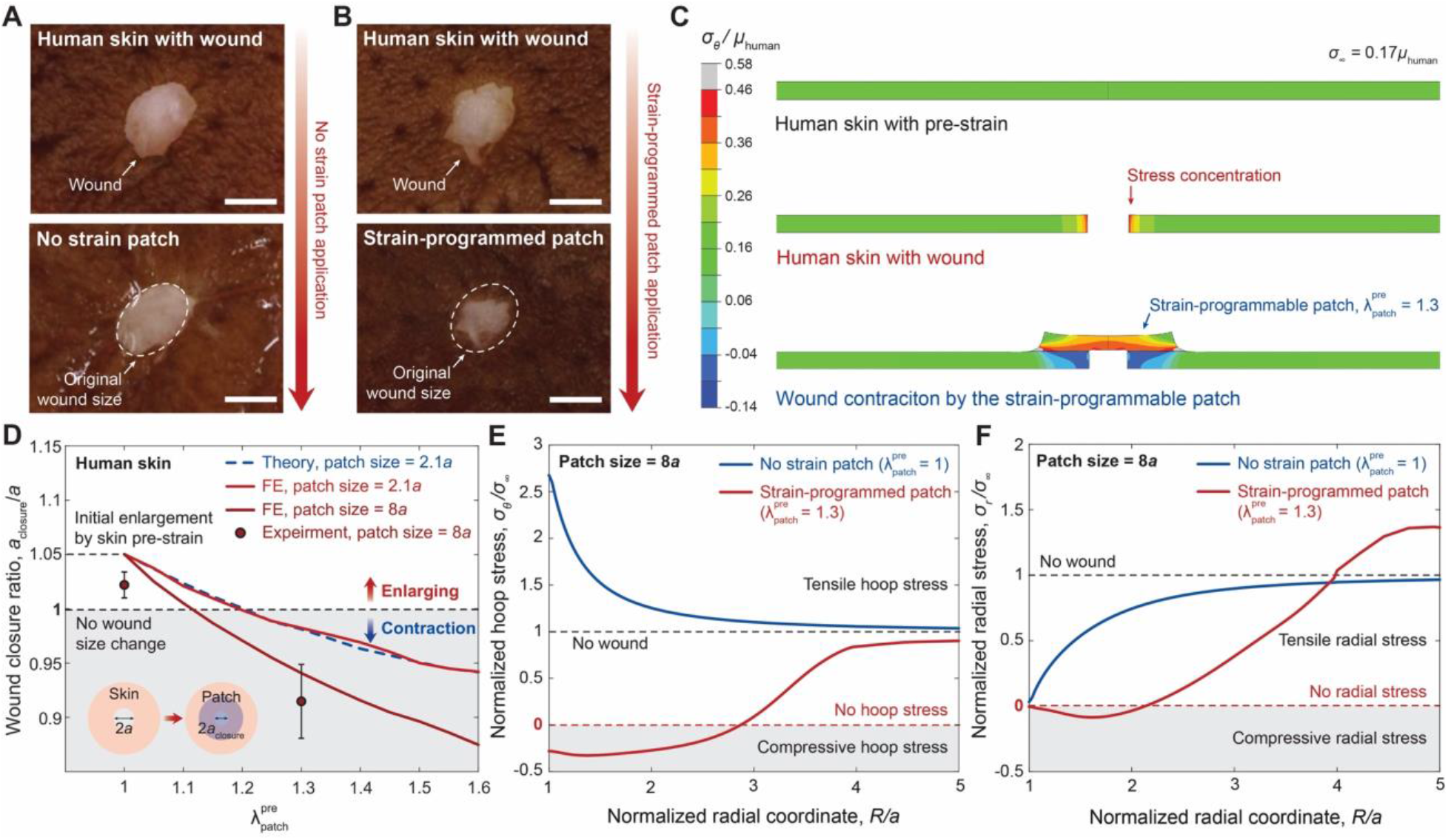
Programmable closure and stress remodeling of *ex vivo* human skin wounds. (**A** and **B**) Images of a no strain patch with 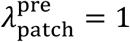 (A) and a strain-programmed patch with 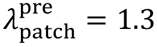 (B) on *ex vivo* human skin with a circular wound. (**C**) Representative finite-element results of the human skin with a strain-programmable patch with 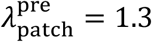 and patch diameter 4 times of the wound diameter. The shear modulus of the human skin is denoted as *µ*_human_, the hoop stress in the human skin as *σ*_θ_, and the residual stress in the intact human skin as *σ*_∞_. (**D**) Finite-element and experimental results for the wound closure ratio as a function of 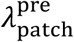. FE, finite-element. (**E** and **F**) Finite-element results for the hoop *σ*_θ_ (E) and the radial *σ*_r_ (F) stresses around the wound with the strain-programmed patches with varying 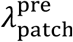. Values in D represent the means ± SD (*n* = 3). Scale bars, 5 mm (A and B).

**Fig. S16.**
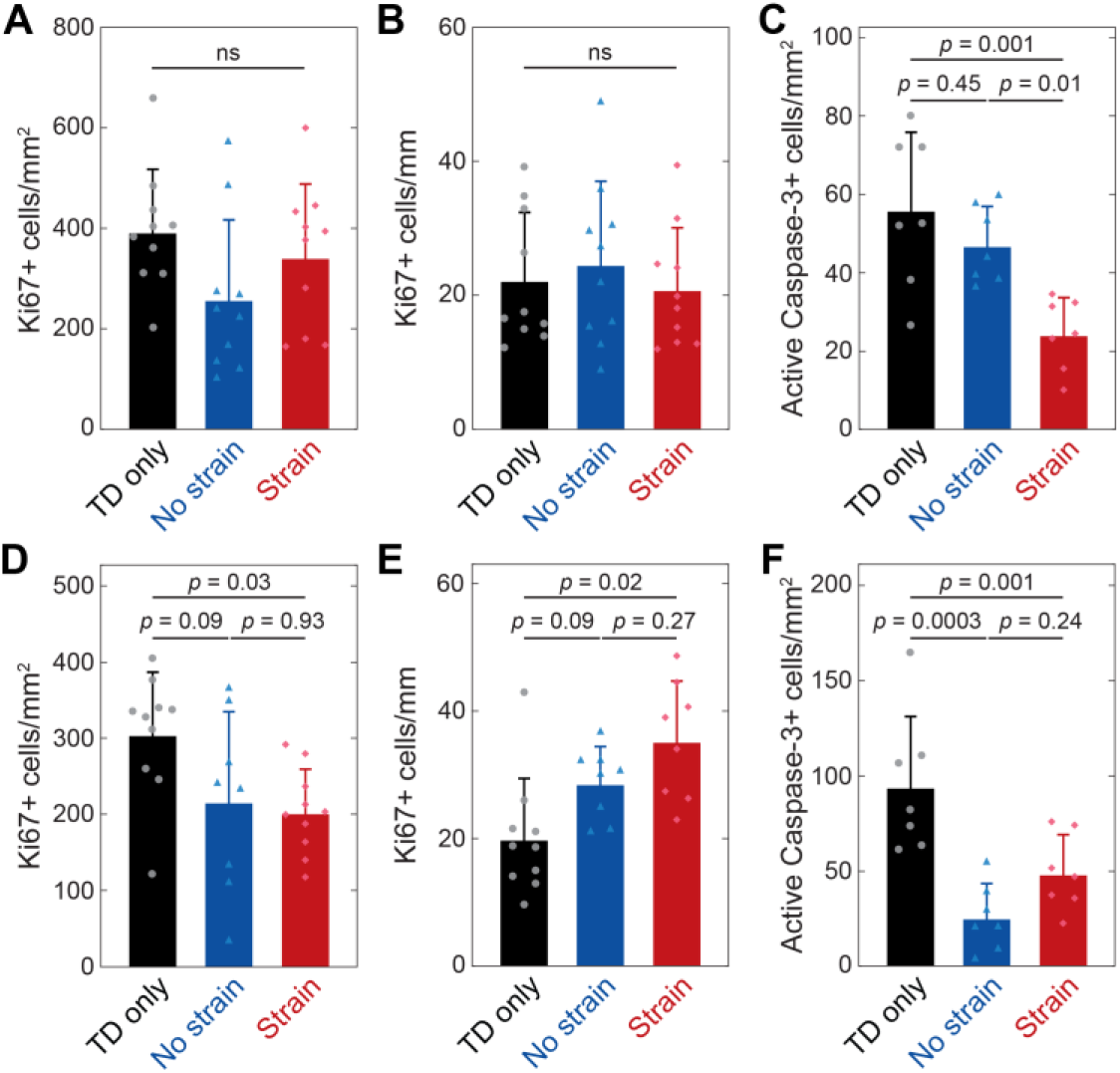
Wound cells proliferation and apoptosis. (**A** and **B**) Quantification of proliferation marker Ki67+ cells in the dermis (A) and the epidermis (B) of day 5 (D5) wounds. (**C**) Quantification of apoptosis marker active Caspase-3+ cells in the dermis of D5 wounds. (**D** and **E**) Quantification of proliferation marker Ki67+ cells in the dermis (D) and the epidermis (E) of day 10 (D10) wounds. (**F**) Quantification of apoptosis marker active Caspase-3+ cells in the dermis of D10 wounds. Values represent the means ± SD (*n* = 7-10). *P* values were derived from one way ANOVA with Tukey’s post-hoc tests. ns, not significant.

**Fig. S17.**
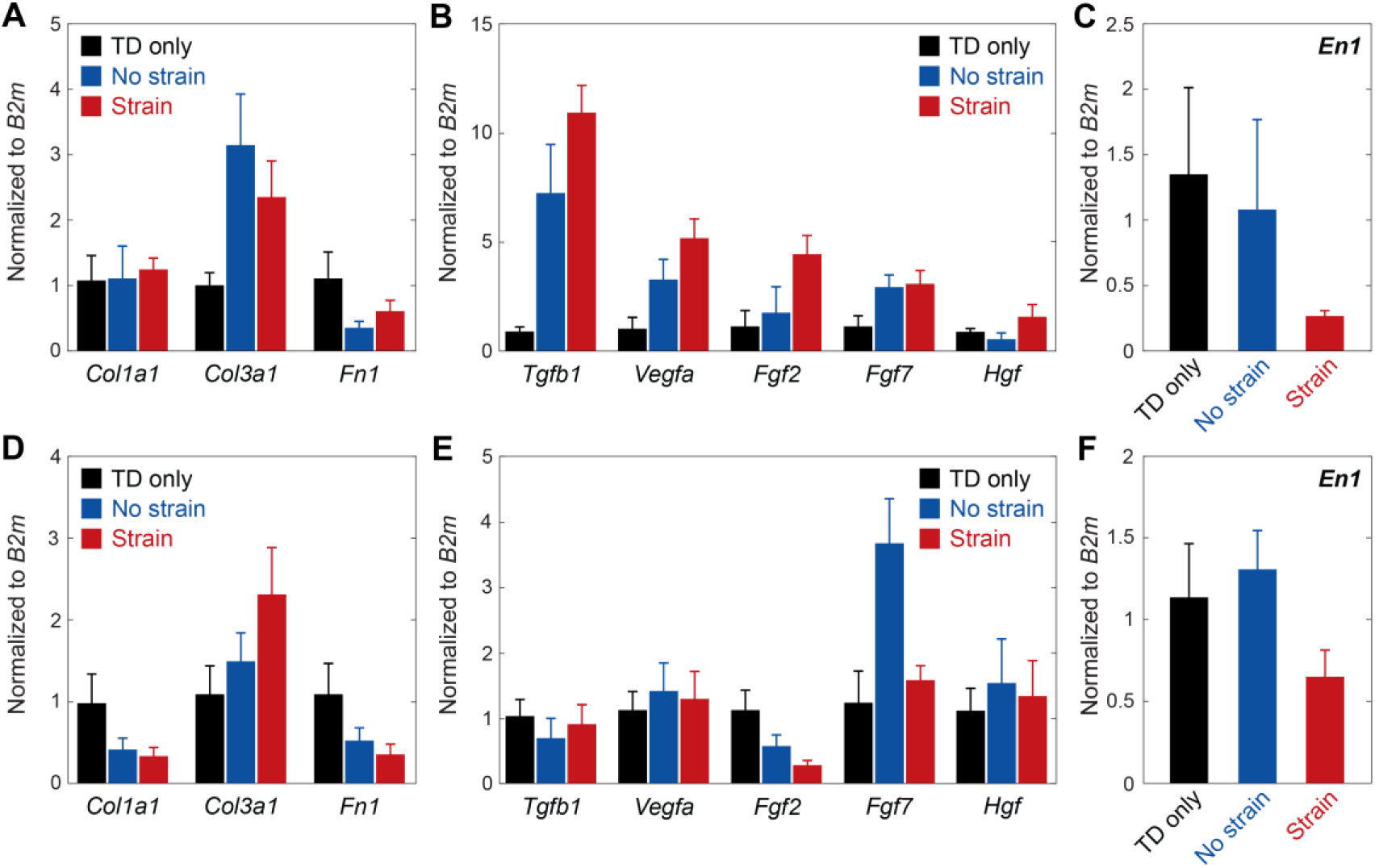
RT-qPCR gene expression analysis. (**A** and **B**) RT-qPCR analysis on day 5 (D5) wounds for ECM-related (A) and growth factor (B) genes. (**C**) *Engrailed-1* (*En1*) wound expression on D5. (**D** and **E**) RT-qPCR analysis on day 10 (D10) wounds for ECM-related (D) and growth factor (E) genes. (**F**) *Engrailed-1* (*En1*) wound expression on D10. Values represent the means ± SEM (*n* = 4-6).

**Fig. S18.**
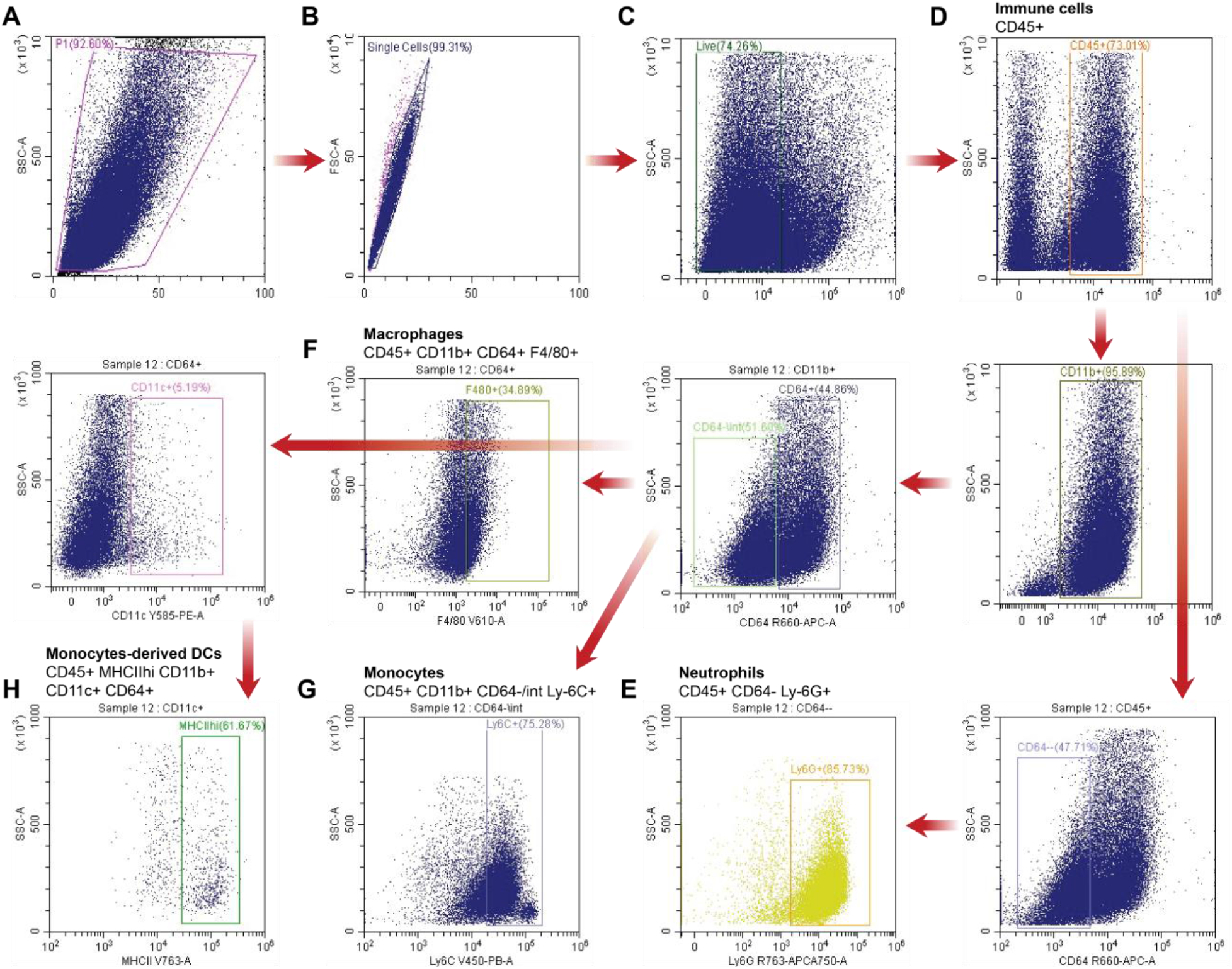
Gating strategy for the flow cytometry analysis of dissociated wound tissues. (**A-H**) Forward and side scatter density plot (A) was used for debris exclusion followed by forward scatter area vs forward scatter height density plot (B) for doublet exclusion. Live (C) immune cells (D) were then characterized as neutrophils (E), macrophages (F), monocytes (G), and monocyte derived dendritic cells (H) with appropriate cell surface antibody staining and sequential gating.

**Fig. S19.**
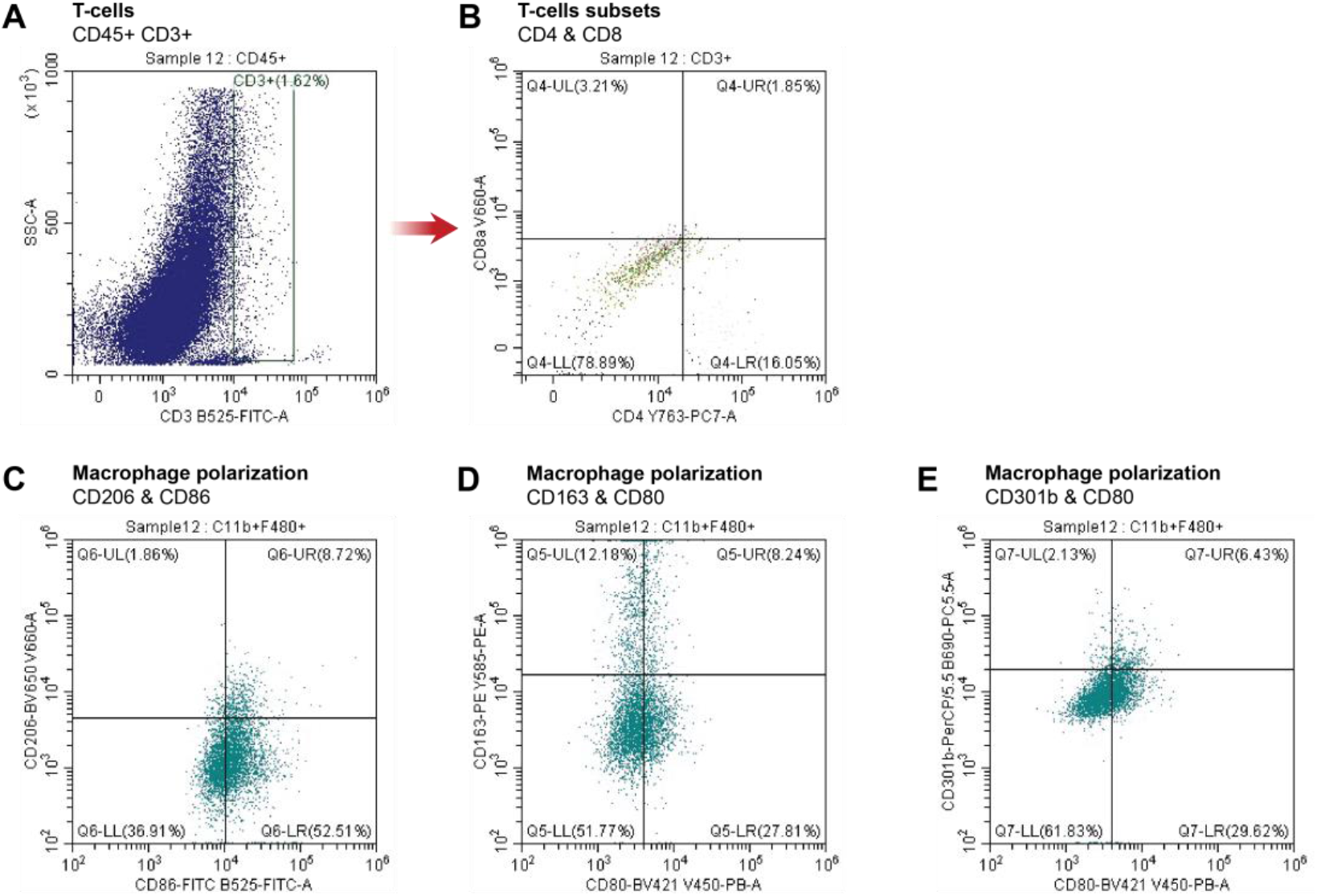
Additional gating strategy for T-cell subsets and polarized macrophages. (**A-E**) Single, live, immune cells were characterized as T-cells (A) and further gated according to CD4 and CD8 expression (B). Density plot graphs were also used to distinguish macrophage (single, live, CD45+CD11b+CD64+F4/80+ cells’) polarized states (C-E).

**Fig. S20.**
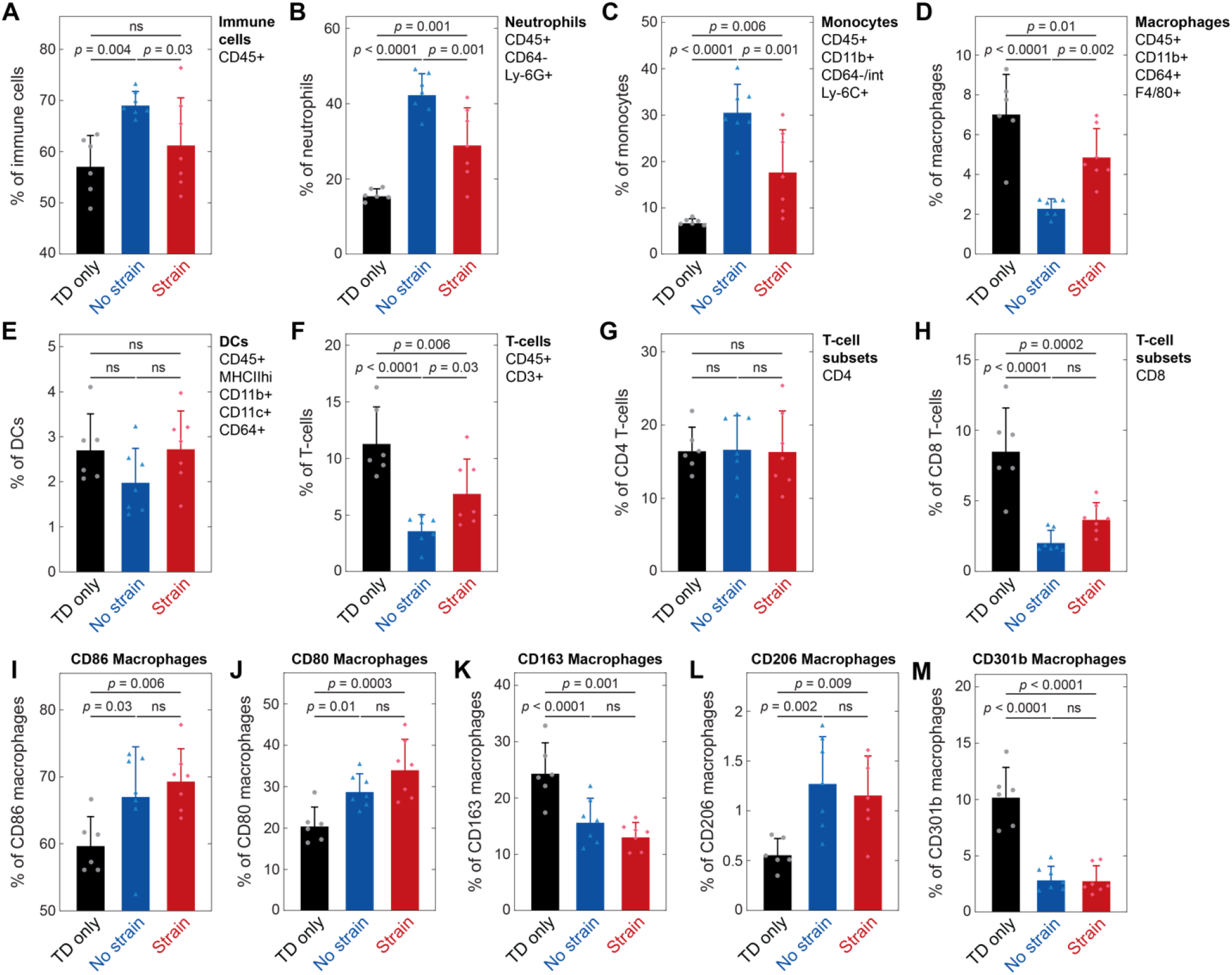
Flow cytometry quantification of major immune cell populations and macrophage polarized states in D5 wounds. (**A-M**) Single cell suspensions were generated from excised wound and peri-wound tissues and stained for the indicated cell surface proteins. Percentage of immune cells (A), neutrophils (B), monocytes (C), macrophages (D), dendritic cells (DCs) (E), and T-cells (F) and T-cell subsets (G and H). Percentage of macrophages expressing markers CD86 (I), CD80 (J), CD163 (K), CD206 (L), and CD301b (M). Each data point represents pooled cells from two mice (four wounds). Values represent the means ± SD (*n* = 6-7). *P* values were derived from one way ANOVA with Fisher’s LSD post-hoc tests. ns, not significant.

**Fig. S21.**
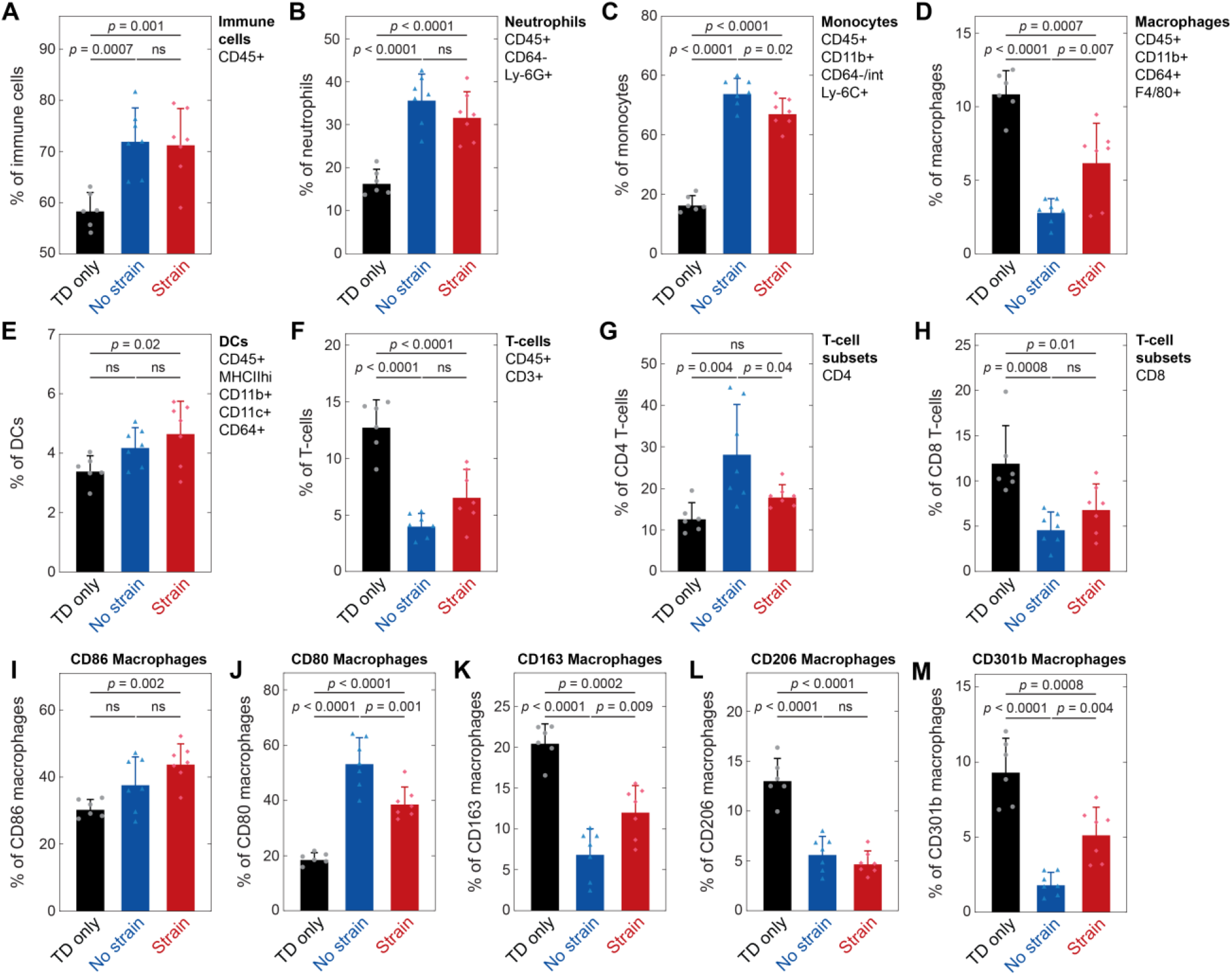
Flow cytometry quantification of major immune cell populations and macrophage polarized states in D10 wounds. (**A-M**) Single cell suspensions were generated from excised wound and peri-wound tissues and stained for the indicated cell surface proteins. Percentage of immune cells (A), neutrophils (B), monocytes (C), macrophages (D), dendritic cells (DCs) (E), and T-cells (F) and T-cell subsets (G and H). Percentage of macrophages expressing markers CD86 (I), CD80 (J), CD163 (K), CD206 (L), and CD301b (M). Each data point represents pooled cells from two mice (four wounds). Values represent the means ± SD (*n* = 6-7). *P* values were derived from one way ANOVA with Fisher’s LSD post-hoc tests. ns, not significant.

**Fig. S22.**
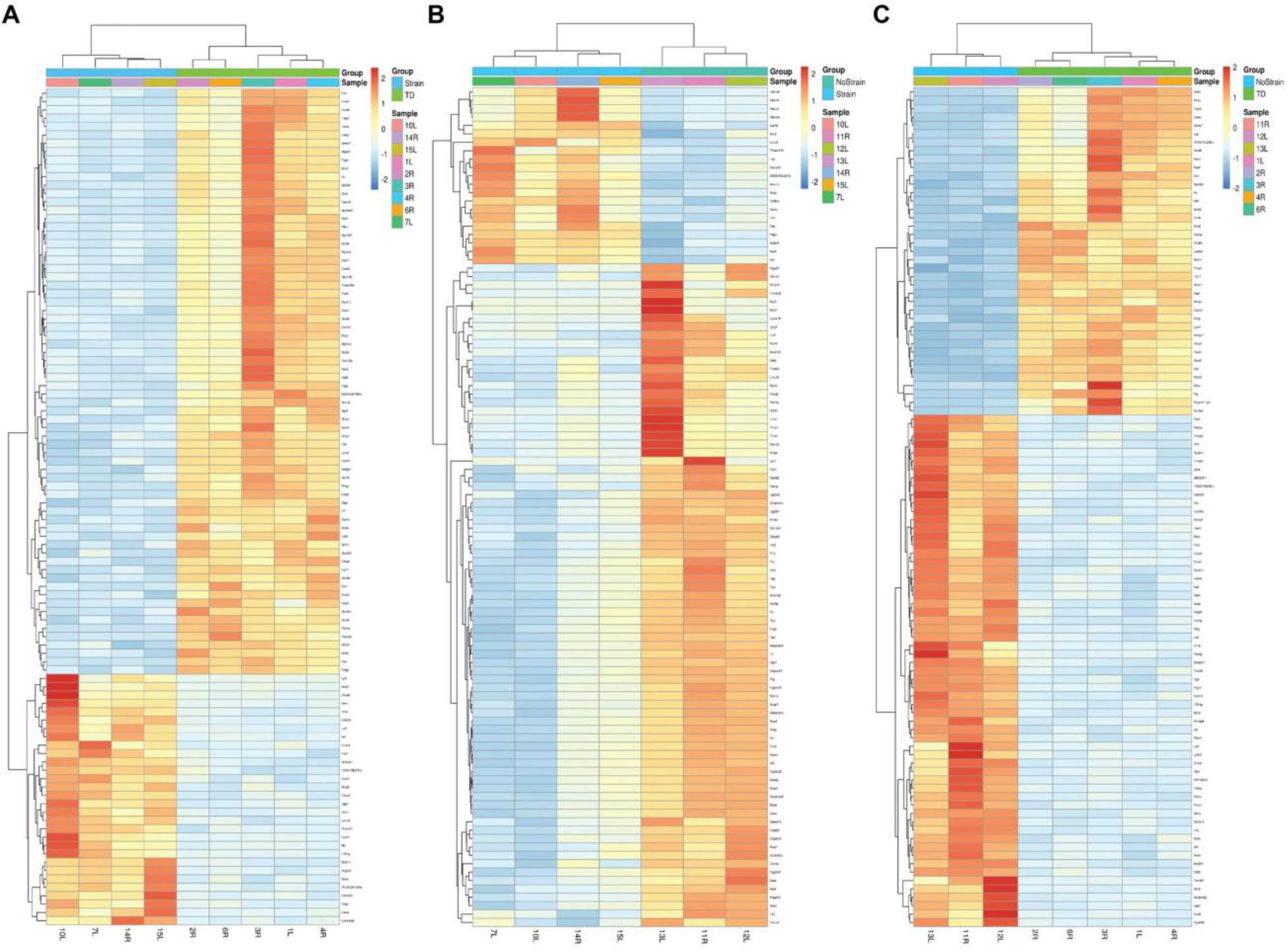
Visualization of RNA sequencing results. (**A-C**) Heatmaps generated from the top 100 differentially expressed features of Strain vs TD (A), No strain vs Strain (B), and No strain vs TD (C) comparisons. Dendrograms were drawn from Ward hierarchical clustering. Higher expression levels correspond to warmer colors.

**Fig. S23.**
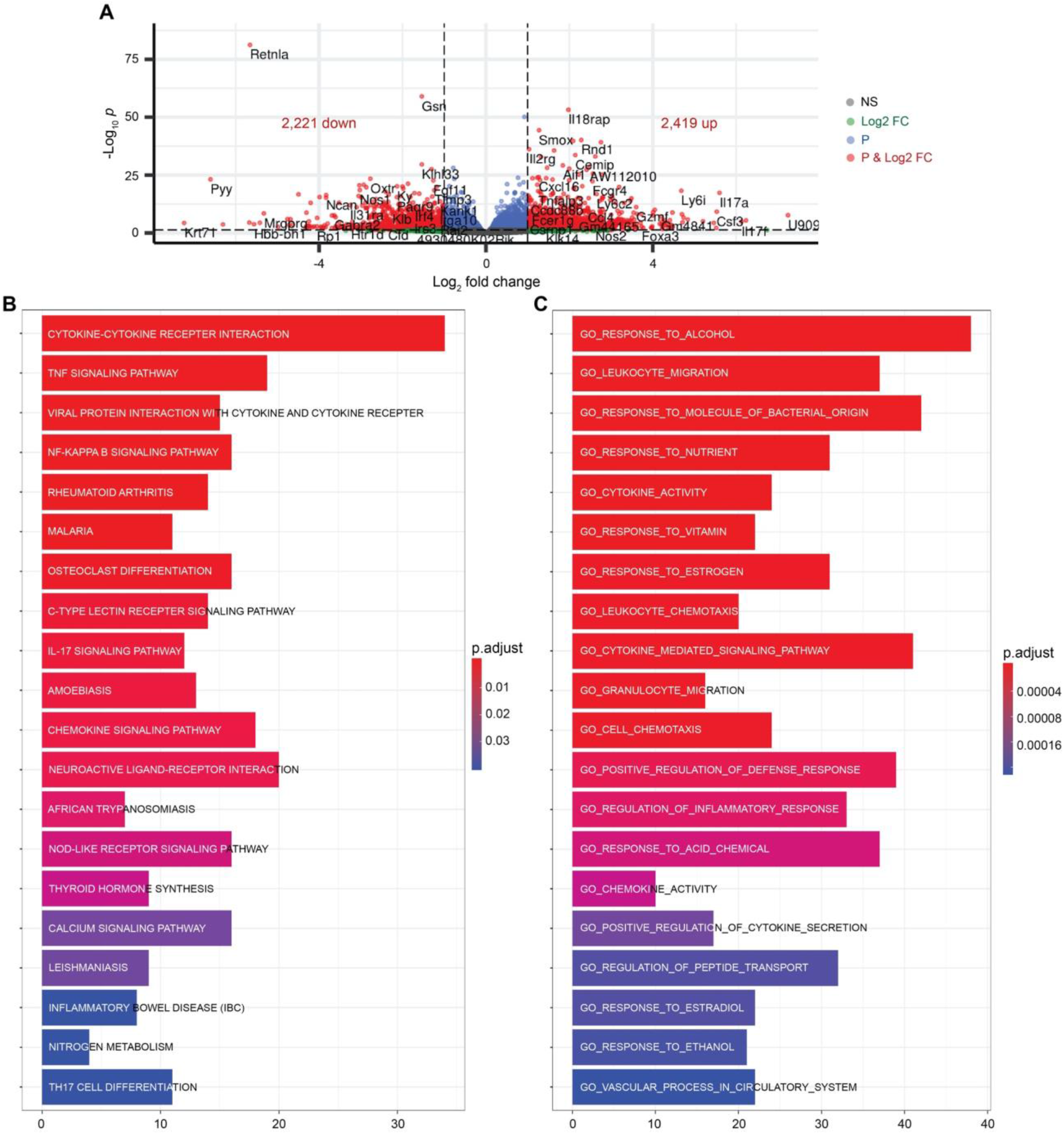
No Strain vs TD differential expression and functional analyses. (**A**) Volcano plot displaying gene expression profile. Red colored data points represent genes that meet the thresholds of fold change (FC) above 1 or under −1, False Discovery Rate (FDR) < 0.05. (**B** and **C**) Functional over-representation analysis utilizing the top 500 differentially expressed genes results in gene ontology (GO) (B) and Reactome (C) databases. The x-axis corresponds to the number of genes implicated in each pathway and the color of the bars correlates with the adjusted *p*-values as shown in the legends.

**Fig. S24.**
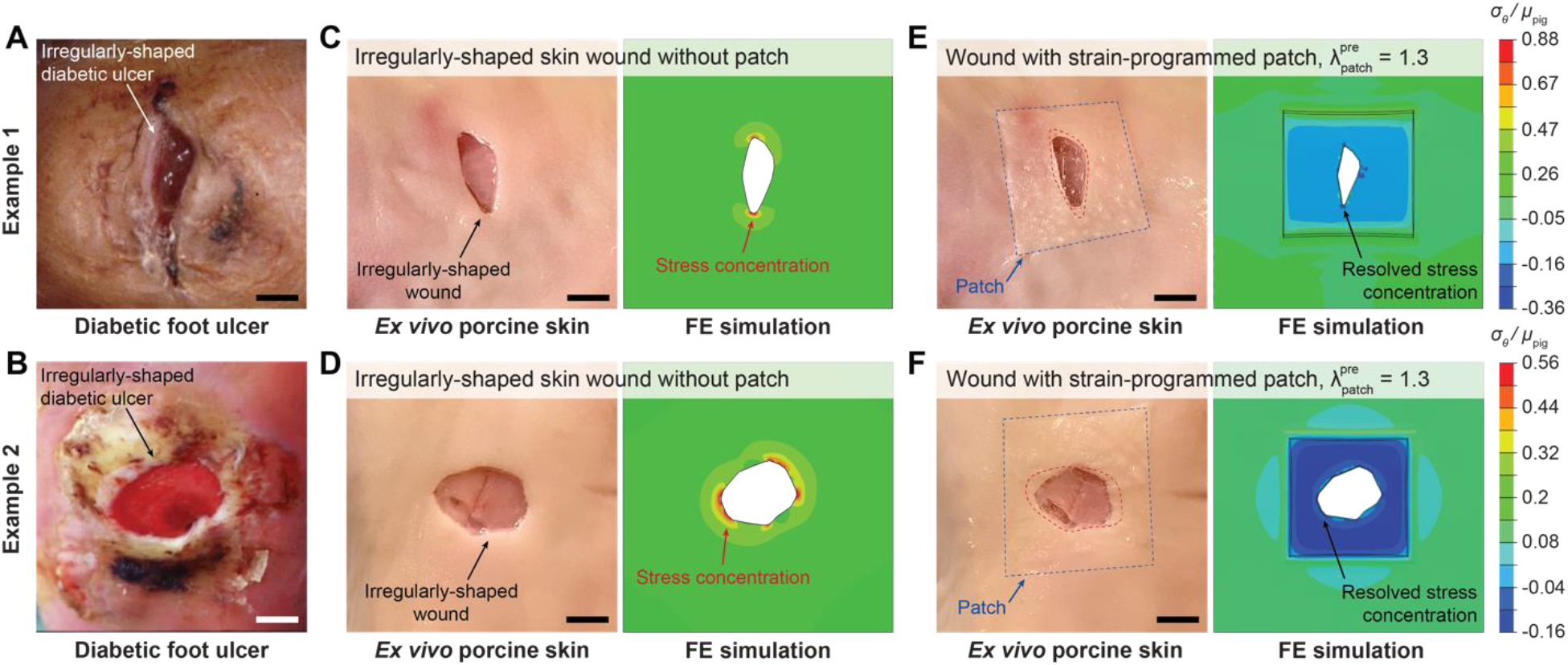
Applicability of the strain-programmable patch to irregularly-shaped wounds. (**A** and **B**) Examples of irregularly-shaped diabetic foot ulcer (DFU) in patients. (**C** and **D**) *Ex vivo* porcine skin wounds and finite-element (FE) model results based on the example DFU. (**E** and **F**) Mechanical modulation of *ex vivo* porcine skin wounds and the corresponding FE model results by the strain-programmed patch 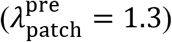. Wound contraction and removal of wound-edge stress concentration are indicated in the experimental images and FE results. The shear modulus of the porcine skin is denoted as *µ*_pig_ and the hoop stress in the porcine skin as *σ*_θ_. Scale bars, 5 mm (A-F).

**Fig. S25.**
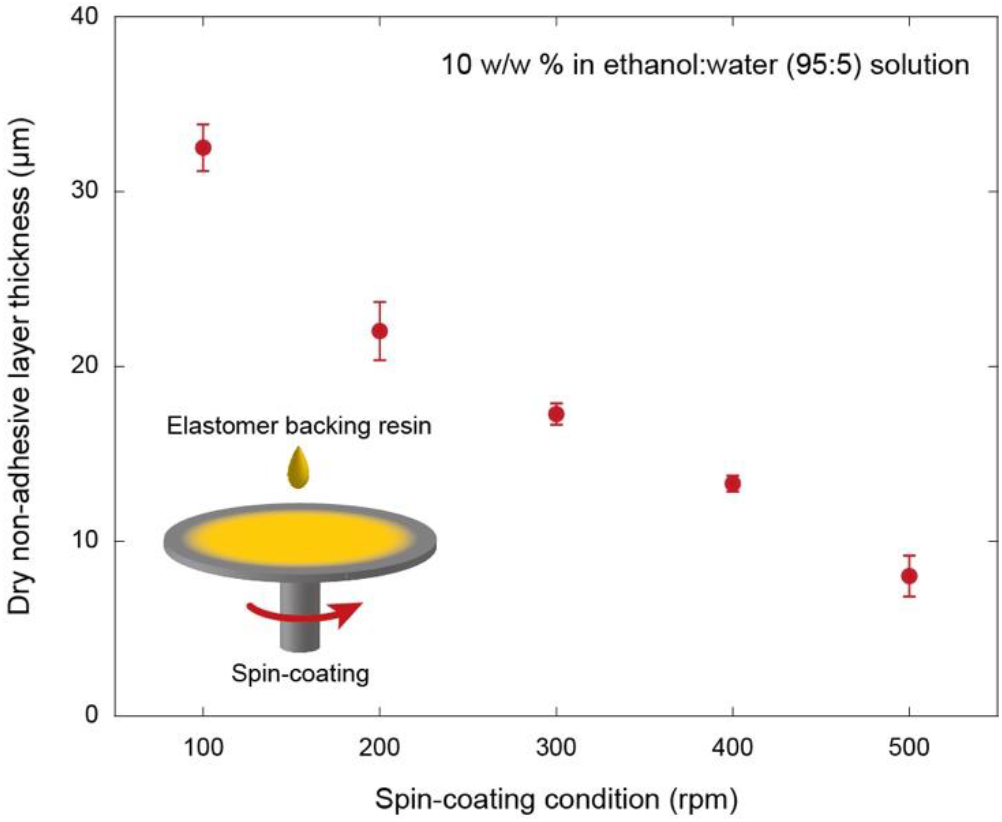
Spin-coating of backing layer. Values represent the means ± SD (*n* = 4).

**Fig. S26.**
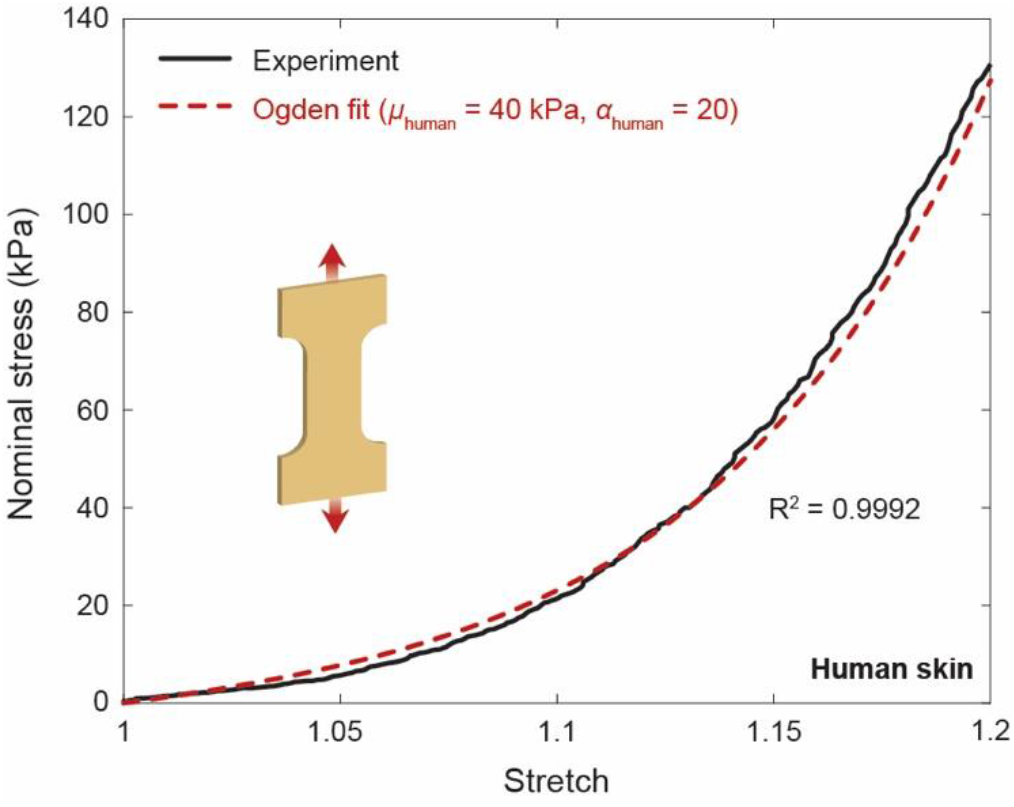
Tensile properties of *ex vivo* human skin. Nominal stress vs. stretch curve for an *ex vivo* human skin fitted with the incompressible Ogden hyperelastic model.

## Captions for other supplementary materials

**Data file S1**. Complete lists of RNA-seq data.

**Data file S2**. Complete list of the antibody cocktail for flow cytometry.

**Movie S1**. Rapid adhesion and closure of an *ex vivo* porcine skin wound by an isotropically strain-programmed patch.

**Movie S2**. Rapid adhesion and closure of an *ex vivo* porcine skin wound by an anisotropically strain-programmed patch.

**Movie S3**. On-demand removal of an adhered patch on an *ex vivo* porcine skin wound by applying a triggering solution.

**Movie S4**. Rapid adhesion and closure of wound in an *ex vivo* diabetic mouse skin by a strain-programmable patch.

## Notes

### Competing Interest Statement

The authors have declared no competing interest.

